# An immunome perturbation is present in juvenile idiopathic arthritis patients who are in remission and will relapse upon anti-TNFα withdrawal

**DOI:** 10.1101/656124

**Authors:** Jing Yao Leong, Phyllis Chen, Joo Guan Yeo, Fauziah Ally, Camillus Chua, Sharifah Nur Hazirah, Su Li Poh, Lu Pan, Liyun Lai, Elene Seck Choon Lee, Loshinidevi D/O Thana Bathi, PRCSG, Thaschawee Arkachaisri, Daniel J. Lovell, Salvatore Albani

## Abstract

**Objectives:** Biologics treatment with anti-TNFα is efficacious in juvenile idiopathic arthritic (JIA) patients. Despite displaying clinical inactivity during treatment, many patients will flare upon cessation of therapy. The inability to definitively discriminate patients who will relapse or continue to remain in remission after therapy withdrawal is currently a major unmet medical need. CD4 T cells have been implicated in active disease, yet how they contribute to disease persistence despite treatment is unknown.

**Methods:** We interrogated the circulatory reservoir of CD4^+^ immune subsets at the single cell resolution with mass cytometry (CyToF) of JIA patients (n=20) who displayed continuous clinical inactivity for at least 6 months with anti-TNFα, and were subsequently withdrawn from therapy for 8 months, and scored as relapse or remission. These patients were examined prior to therapy withdrawal for putative subsets that could discriminate relapse from remission. We verified on a separate JIA cohort (n=16), the continued dysregulation of these circulatory subsets 8 months into therapy withdrawal. The immunological transcriptomic signature of CD4 memory in relapse/remission patients was examined with Nanostring.

**Results:** An inflammatory memory subset of CD3^+^CD4^+^CD45RA^−^TNFα^+^ T cells deficient in immune checkpoints (PD1^−^CD152^−^) was present in relapse patients prior to therapy withdrawal. Transcriptomic profiling reveals divergence between relapse and remission patients in disease centric pathways involving (a) TCR activation, (b) apoptosis, (c) TNFα, (d) NF-kB and (e) MAPK signalling.

**Conclusions:** A unique discriminatory immunomic and transcriptomic signature is associated with relapse patients and may explain how relapse occurs.

## INTRODUCTION

Targeted therapy of juvenile idiopathic arthritis (JIA) with anti-TNFα biologics is efficacious with 70-80% responders and up to 50% achieving clinical inactivity on long term treatment[1, 2]. While sustained immune suppression through anti-TNFα is generally well tolerated, clinicians seek to achieve clinical remission off medication, to reduce risk of general infection, adverse events and financial burden[2]. Drug withdrawal in patients who attain clinical inactivity is complicated by the fact that 50-80% patients relapse upon therapy discontinuation[3, 4]. This phenomenon indicates that relapse patients who have attain clinical inactivity on medication, as defined by Wallace criteria, continue to experience subclinical inflammation and persistence of disease without overt presentation of clinical symptoms. Conversely, patients who achieve clinical remission off medication could be spared long-term drug effects. As such, there is a clinical need to address how discontinuing anti-TNFα therapy can be safely implemented, and a scientific need to understand the immune mechanisms related to relapse.

The etiology of JIA remains widely debated[5]. Unsupervised genome wide association studies and pathway analysis have highlighted the role of CD4 T-helper cell populations in autoimmune disease progression[6]. CD4 T cells were shown to infiltrate the synovium micro-environment[7–10], and corresponding pathogenic CD4 HLA-DR^+^ T-effector and regulatory subsets possessing strong immune phenotypic and TCR correlation with synovial T cells have been found re-circulating in the blood during active inflammation[11, 12]. Furthermore, epigenetic histone modifications associated with enhancer functions has been detected in CD4 T cells of JIA patients[13]. Using network analysis of DNA CpG methylation sites in total CD4 T cells, we have previously demonstrated that T-cell activation pathways are associated with clinical fate upon anti-TNFα withdrawal [14]. However, the identity of the specific pathogenic CD4 subset that maintains subclinical disease persistence remains elusive.

We have leveraged on the high dimensional single cell resolution capabilities of Cytometry by Time-of-Flight (CyToF) to uncover the CD4 T cell subset responsible for disease persistence. JIA patients who maintained clinical inactivity under anti-TNFα for at least 6 months, were subsequently withdrawn from therapy. These patients were rigorously scored for their disease progression across 8 months and their clinical outcome defined as relapse or remission. We interrogated the circulatory immunome of JIA patients prior to therapy withdrawal, and identified an inflammatory memory CD3^+^CD4^+^CD45RA^−^TNFα^+^ T cell that is deficient for immune checkpoints (PD1^−^CD152^−^) in individuals who will relapse. Separately in patients who have flared after 8 months, this memory subset has extended towards TNFα^+^IL-6^+^. We were able to use the memory CD3^+^CD4^+^CD45RA^−^TNFα^+^ subset to discriminate relapse from remission patients prior to withdrawal, and cross validated this dysregulation against a larger cohort of healthy paediatric controls. Finally, transcriptomic profiling of CD4 memory cells reveal disease centric pathways that reflect selective divergence among relapse and remission patients. This study underscores immunological differences at the core of the dichotomic clinical responses. These differences provide a framework for understanding the mechanisms of disease relapse upon drug withdrawal, thus providing a knowledge-based guidance for management while proposing some potential new targets for intervention.

## MATERIALS AND METHODS

### Samples

Peripheral blood mononuclear cells (PBMCs) were obtained from polyarticular JIA patients recruited through the “*Determining Predictors of Safe Discontinuation of Anti-TNF treatment in JIA*” trial (ID: NCT00792233)[15]. Patients were treated with anti-TNFα biologics and shown to be in an inactive disease state for 6 months, were enrolled into the study. Clinical inactivity was defined by Wallace criteria[16]: (a) absence of active joints, (b) lack of fever, rash, serositis attributable to JIA, (c) no active uveitis, (d) within normal range of ESR unless attributable to JIA, (e) physician global disease activity of ≤ 0.5 Likert-like scale and (f) duration of morning stiffness ≤ 15 minutes. Upon enrollment, patients are withdrawn from anti-TNFα therapy and accessed through monthly clinical visits for a study period of 8 months. Clinical outcome is designated as relapse or remission depending on six core JIA parameters; (a) number of active joints, (b) number of joints with loss of motion, (c) medical doctor global assessment of current disease activity (Likert-like scale), (d) patient/parent global assessment of overall disease severity in prior week (Likert-like scale), (e) a validated measure of physical function (CHAQ) and (f) ESR. A patient was considered to be experiencing a relapse if there was ≥ 30% worsening in more than three of the six JIA core parameters, and with no more than one parameter improving by > 30%. For remission individuals, they would have achieved ≥ 14 months of clinical inactivity from prior recruitment to study end. PBMCs were interrogated by CyToF from patients (n=20) prior to withdrawal were designated as (T_o_), and separately from another batch (n=16) at the end of 8 months after withdrawal were designated as (T_end_). Patient PBMCs (n=12) were also sorted for CD3^+^CD4^+^CD45RO^+^CD45RA^−^ for Nanostring analysis. The demographics/medication history profile of JIA patients withdrawn from therapy and sample usage breakdown is as shown (**online supplementary Table S1**).

Age-matched healthy controls (n=69) were recruited through the Precision Rheumatology International Platform (PRIP) study conducted at the KK Women’s and Children’s Hospital (KKH). These controls have no indication of inflammation and PBMCs were isolated pre-operatively from patients scheduled for day surgeries. Healthy PBMCs were examined with CyToF (n=10), Nanostrong (n=3) or age-matched strata cross validation for ROC curve (n=56).

Paired treatment naïve/post treatment JIA patients (n=4) were also recruited through the study *“A precision medicine approach to understand and predict responsiveness to therapy in human arthritis*” conducted in KKH for Nanostring analysis. These patients with active JIA were initially treatment naive to anti-TNFα, and after a 6-month drug course exhibited treatment susceptibility determined by complete absence of active joints. The demographics/medication history profile of JIA patients is as shown (**online supplementary Table S2**).

Additional methodological details are available as supplementary information.

## RESULTS

### CD4^+^CD45RA^−^TNFα^+^ T cells is present in JIA patients prior to relapse

Dsyregulated CD4 T cells are thought to contribute to JIA pathogenesis[7–12]. We interrogated the circulatory CD4 landscape of JIA patients (n=20) prior to therapy withdrawal to understand why certain individuals relapse. At this stage, the patients are clinically scored to be inactive for 6 months, thus patients who will relapse or remain in remission are clinically indistinguishable prior to withdrawal. We assessed the PBMCs with a CyToF panel consisting of 31 functional, 6 lineage markers (**online supplementary Table S3**) and CD45 barcoding to facilitate pooling of individuals[17]. Batch variability in staining was monitored through an internal biological control (**online supplementary Figure S1**). The debarcoded CD3^+^CD4^+^ T cells were exported, normalized for cell events and the 31 markers dimensionally reduced with MarVis onto a bivariate X-Y axis through t-SNE (**online supplementary Figure S2**). Clustering with k-means segregated the CD4 cells into distinct nodes (**Figure 1A**), and we detected an enrichment (p < 0.01) in node 22 for patients who will relapse, that represents CD4^+^CD45RA^−^TNFα^+^IFNγ^−^CD152^−^PD1^−^ T cells (**Figure 1A-C**). To ensure the results are not due to clustering artefacts, we manually gated the pre-clustering FCS files (**Figure 1D**, gating strategy in **online supplementary Figure S3**), and determined significant (p < 0.05) up-regulation in CD4^+^CD45RA^−^ memory subsets that was restricted within the TNFα^+^ compartment. In particular, relapse patients were enriched for CD4^+^CD45RA^−^TNFα^+^ T cells which were absent for IFNγ expression, and were notably deficient in immune checkpoints (PD1/CD152). We investigated the relationship of the dysregulated T effectors and immune checkpoint expression within the memory compartment (**Figure 1E**). There was a stronger positive correlation of CD45RA^−^TNFα^+^ with CD45RA^−^CD152^−^PD1^−^ (r = 0.8257) as opposed to CD45RA^−^CD152^+^PD1^+^ (r = 0.5987) cells across the patients. The percentage of TNFα^+^ cells was significantly higher (p < 0.0001) in CD45RA^−^CD152^−^/PD1^−^ cells (**Figure 1F**). While relapse patients showed a perceptible increase in T-effectors in the absence CD152/PD1, there was a marked drop to up-regulate CD152/PD1 as compared with remission patients (**Figure 1G-H**).

**Figure 1.**
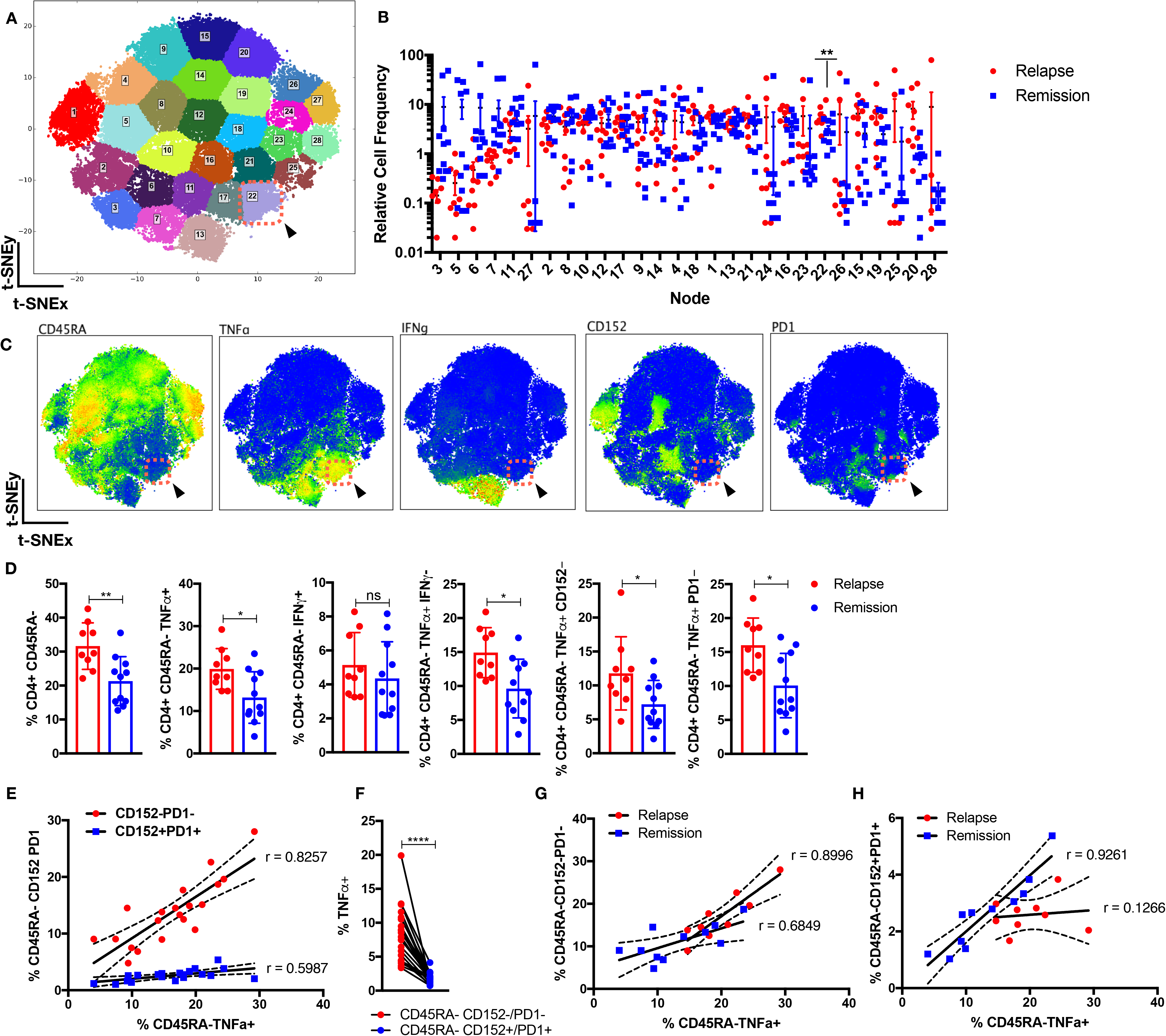
Perturbation in CD4 landscape in JIA patients who will relapse. Circulatory CD3^+^CD4^+^ cells from JIA patients (n=20; relapse =9, remission =11) prior to therapy withdrawal were stained with 31 functional markers in CyToF and dimensionally reduced onto a bivariate X-Y axis scale with t-SNE. (**A**) The t-SNE map is segregated into 28 distinct nodes with k-means, and node 22 is highlighted (red-dotted box). (**B**) The distribution of relative cell frequency in relapse or remission patients across the nodes is shown, with node 22 significantly higher (p < 0.01) in relapse individuals. (**C**) The phenotypic expression of markers is shown for node 22 (red-dotted box). (**D**) Supervised gating of the pre-clustering FCS files validates the relevant CD4 memory cellular subsets in relapse versus remission individuals. (**E**) Correlation analysis of CD45RA^−^TNFα^+^ versus CD45RA^−^CD152^−^/PD1^−^ or CD45RA^−^CD152^+^/PD1^+^ as percentage of CD3^+^CD4^+^ cells across JIA patients. (**F**) Percentage of TNFα^+^ cells in CD45RA^−^CD152^−^/PD1^−^ or CD45RA^−^CD152^+^/PD1^+^ compartment. Correlation analysis of CD45RA^−^TNFα^+^ versus (**G**) CD45RA^−^CD152^−^/PD1^−^ or (**H**) CD45RA^−^ CD152^+^/PD1^+^ as percentage of CD3^+^CD4^+^ cells across relapse or remission patients. Comparison of cellular subsets performed with Mann-Whitney U, unpaired or paired two tail test, means ± S.D., * p < 0.05, ** p < 0.01. Correlation analysis performed with Pearson correlation, two tail test.

### CD4^+^TNFα^+^ healthy landscape unveils subclinical T-effector diversification in relapse patients

To validate the findings above and to investigate for the possibility of disease centric CD4^+^ cellular subsets that are masked by comparing JIA relapse/remission individuals, we included age-matched paediatric healthy controls. CD4^+^TNFα^+^ T cells from JIA relapse/remission (n=20) prior to withdrawal or healthy individuals (n=10) were compared to further delineate key differences within the T effector compartment. JIA patients who will relapse were enriched (p < 0.05) for node 17 against remission individuals (**Figure 2A-B**), and additionally for nodes 17,13 and 18 against healthy controls (p <0.05) (**Figure 2C-D**). Node 17 reaffirms relapse individuals are enriched for CD4^+^CD45RA^−^TNFα^+^IFNγ^−^CD152^−^PD1^−^ T cells (**Figure 2E-F**) as compared with remission or healthy individuals. Additionally node 13 (**Figure 2E**) represents CD4^+^CD45RA^−^TNFα^+^IFNγ^−^CD152^−^PD1^−^ T cells that are also IL-6^+^. Supervised gating (**Figure 2G** gating strategy in **online supplementary Figure S3**) validates that relapse individuals are enriched (p < 0.001) with CD4^+^CD45RA^−^TNFα^+^IL-6^+^ as compared with healthy individuals. The intensity of TNFα is significantly higher (p < 0.0001) in CD45RA^−^IL-6^+^ as opposed to IL-6^−^ cells (**Figure 2H**), and relapse individuals has higher fold increase (p < 0.05) in TNFα in CD45RA^−^IL-6^+^ cells as compared with remission or healthy individuals.

**Figure 2.**
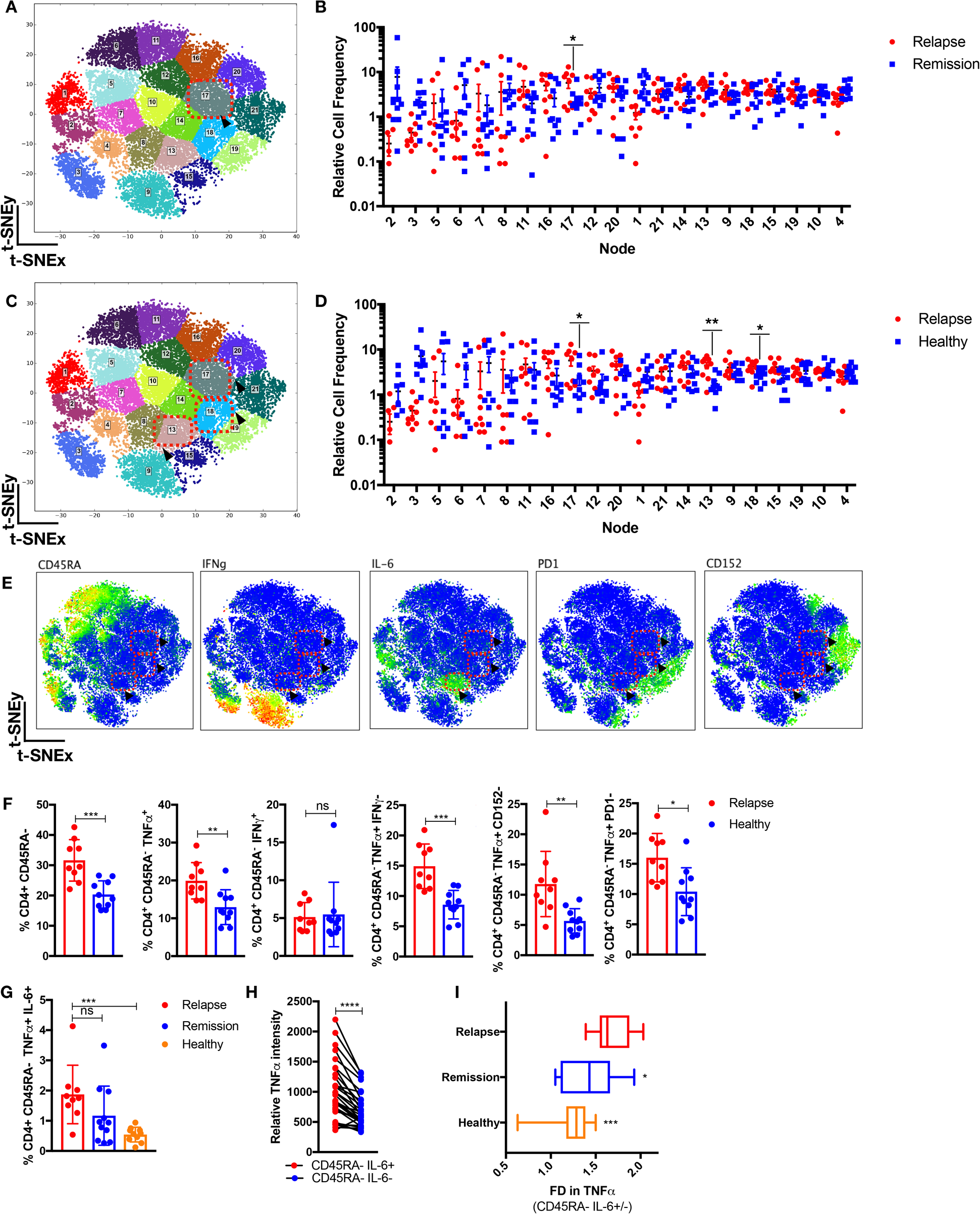
Subclinical T-effector diversification in relapse individuals. CD4^+^TNFα^+^ cells from JIA patients (n=20; relapse =9, remission =11) prior to therapy withdrawal and healthy paediatric controls (n=10) were analysed through t-SNE. (**A**) The CD4^+^TNFα^+^ t-SNE map is segregated into 21 nodes with enrichment in (**B**) node 17 (p < 0.05) for relapse as compared with remission individuals and (**C-D**) node 17,18 and 13 (p <0.05) for relapse as compared with healthy individuals. (**E**) The phenotype expression is shown for node 17,18 and 13 (red-dotted boxes). (**F**) Supervised gating of pre-clustering FCS files validates the relevant CD4 memory cellular subsets in relapse versus healthy individuals. (**G**) Supervised gating of CD4^+^CD45RA^−^TNFα^+^IL-6^+^ cells in relapse/remission/healthy individuals. (**H**) Relative TNFα intensity in CD45RA^−^IL-6^+^ or CD45RA^−^IL-6^−^ cells. (**I**) The fold difference in TNFα intensity in CD45RA^−^IL-6^−^ over CD45RA^−^IL-6^+^ cells in relapse/remission/healthy individuals. Comparison of cellular subsets performed with Mann-Whitney U, unpaired or paired two tail test, means ± S.D., * p < 0.05, ** p < 0.01.

### Overt T-effector diversification during flare manifestation and quiescence in stable remission

We have observed the presence of CD4 memory T cells in patients prior to relapse. To examine this phenomenon further, we interrogated the CD4 landscape of an independent batch of JIA individuals (n=16) withdrawn from therapy for 8 months, that either developed flare or remained in stable remission. Relapse (in flare) patients exhibited the emergence of a previously subclinical CD4^+^CD45RA^−^TNFα^+^IL-6^+^ subset (**Figure 3A-D**, node 7) as compared with patients who remained in remission after 8 months of withdrawal. To determine the state of immunological quiescence in remission (T_o_: prior withdrawal, T_end_: 8 months withdrawal) patients, we gated for the relevant dysregulated CD4^+^CD45RA^−^TNFα^+^ subsets and found no difference as compared with healthy individuals (**Figure 3E**). The CD4 landscape of remission patients (T_o_/T_end_) as compared with healthy individuals revealed mostly similar profiles (**Figure 3F**) except for node 17 and 1 enriched in remission T_o_ or T_end_ patients respectively. Both nodes were absent for TNFα, with node 17 exhibiting CD45RA^+^ and node 1 expressing CD45RA^−/+^CXCR3^+^CCR6^+^ phenotype (**Figure 3G**).

**Figure 3.**
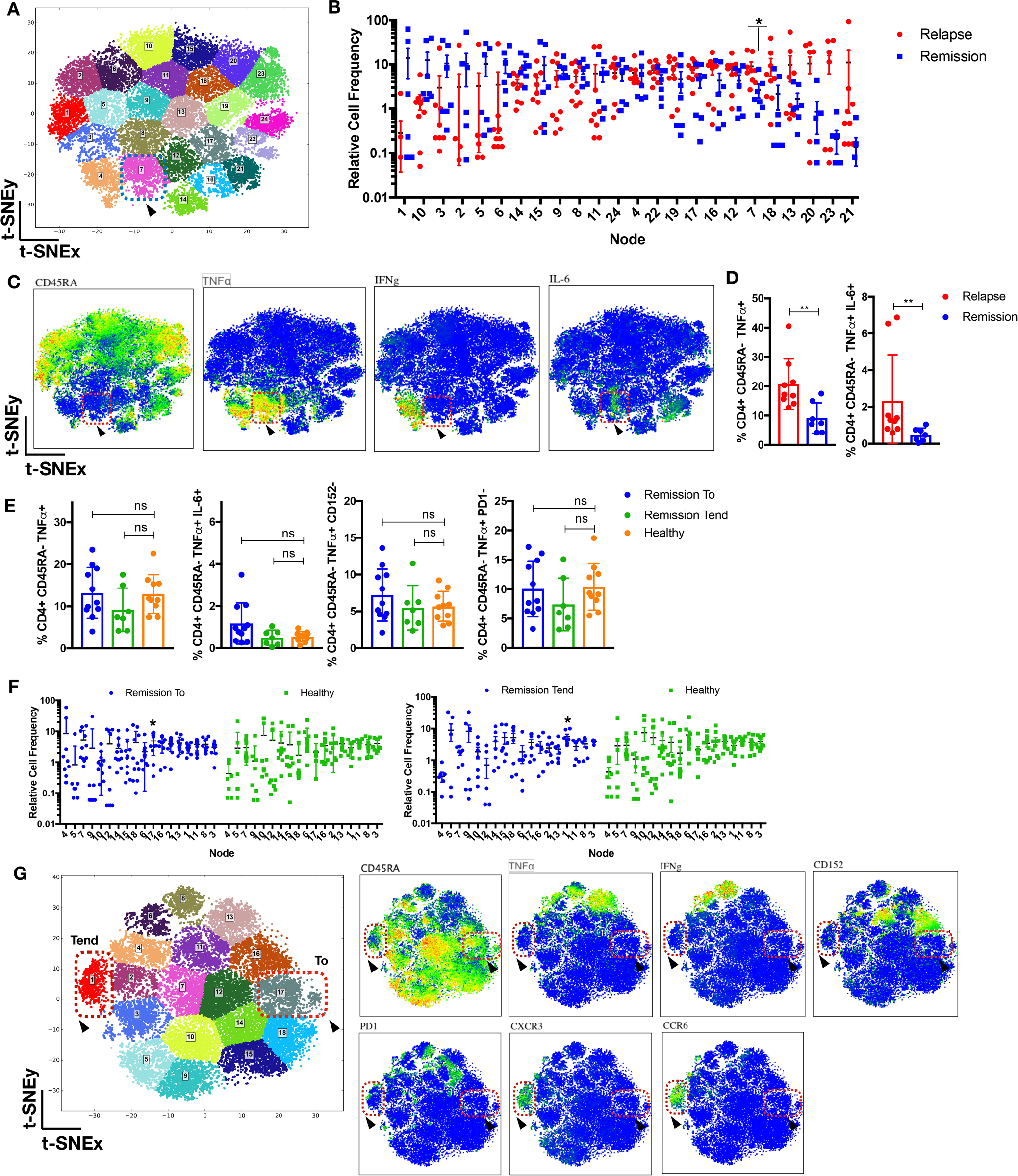
Emergence of T-effector diversification in flare manifestation and quiescence in stable remission. (**A-D**) CD4^+^ T cells from JIA patients (n=16; relapse=9, remission=7) withdrawn from therapy for 8 months were analysed with t-SNE. (**A**) The t-SNE map is segregated into 24 nodes, node 7 is highlighted (blue dotted box). (**B**) The distribution of relative cell frequency across the nodes is shown, with node 7 enriched (p < 0.05) in relapse individuals, exhibiting (**C**) CD4^+^CD45RA^−^TNFα^+^IL-6^+^IFNγ^−^ phenotype (red-dotted box). (**D**) Supervised gating of pre-clustering FCS files for CD4^+^CD45RA^−^TNFα^+^IL-6^+^ in relapse and remission patients. (**E**) Supervised gating of relevant CD4^+^CD45RA^−^TNFα^+^ subsets in remission patients (T_o_: prior to withdrawal or T_end_: 8 months after withdrawal) or healthy individuals. (**F-G**) CD4^+^ T cells from JIA remission patients (n=18; T_o_=11, T_end_=7) and healthy controls (n=10) were analysed with t-SNE. (**F**) The distribution of relative cell frequency across the nodes is shown, with node 17 enriched (p < 0.05) in remission T_o_ and node 1 enriched (p <0.05) in remission T_end_ as compared with healthy individuals. (**G**) Node 1 and 17 phenotypes are depicted. Comparison of cellular subsets performed with Mann-Whitney U, two tail test, means ± S.D., * p < 0.05, ** p < 0.01.

### Corresponding increase of memory Tregs and CD45RA^−^TNFα^+^ prior to relapse

As Tregs have been previously implicated in JIA pathogenesis[10, 12, 18–20], we examined their role in JIA patients prior/after therapy withdrawal (**Figure 4A-B**, gating strategy in **online supplementary Figure S3**). While no differential total Treg frequencies was detected, we observed higher levels of CD45RA^−^Treg (p < 0.0001) in JIA patients prior to relapse. We further verified this with t-SNE analysis of Tregs from relapse/remission patients prior to withdrawal, and determined an enrichment for CD45RA^−^CD152^+^CD127^−^Tregs for relapse individuals (**Figure 4C-D**). Whilst no correlation (r=0.1532) with total Tregs was observed for CD45RA^−^TNFα^+^ cells (**Figure 4E**), there was a positive correlation (r=0.6017) with CD45RA^−^Treg.

**Figure 4.**
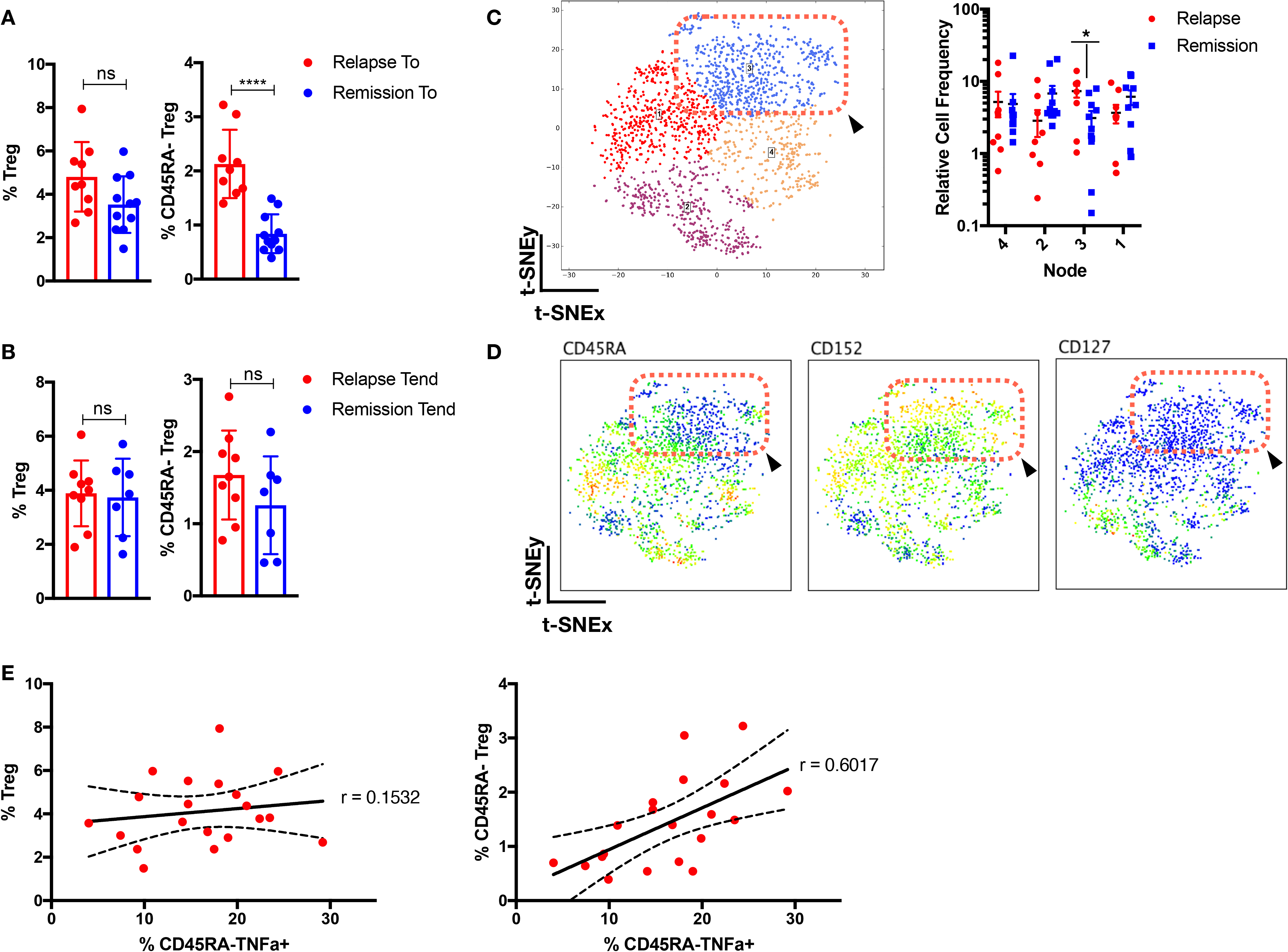
Corresponding increase of memory Tregs and CD45RA^−^TNFα^+^ prior to relapse. Supervised gating of Tregs and CD45RA^−^Tregs in JIA patients (**A**) prior (n=20) and (**B**) after withdrawal (n=16) of therapy. (C-D) Tregs cells from JIA patients (n=20; relapse=9, remission=11) prior to withdrawal is analysed with t-SNE and (**C**) segregated into 4 nodes, with node 3 (p <0.05) enriched in relapse individuals. (**D**) Phenotype of node 3 is depicted. (E) Correlation analysis of frequency of Tregs or CD45RA^−^Tregs with CD45RA^−^TNFα^+^. Comparison of cellular subsets performed with Mann-Whitney U, two tail test, means ± S.D., * p < 0.05, **** p < 0.0001. Correlation analysis performed with Pearson correlation, two tail test.

### Transcriptomic divergence in disease centric pathways that persists despite therapy

We have previously shown that JIA patients who developed active disease upon therapy withdrawal have stable epigenetic DNA CpG modifications in CD4 T cells that predisposed towards T cell activation and TCR signaling[14]. We wanted to investigate if CD4 memory T cells from JIA relapse/remission patients were differential in transcriptomic profile when their TCR is activated. We sorted for CD3^+^CD4^+^CD45RA^−^CD45RO^+^ T cells (**online supplementary Figure S4**), stimulated 24hrs with anti-CD3/CD28, and profiled 579 immunological genes through Nanostring. Functional gene enrichment analysis (DAVID) of JIA relapse (n=6) or remission (n=6) patients versus healthy individuals (n=3) identified 5 common disease centric pathways that persisted from prior to after withdrawal of therapy (**online supplementary Figure S5 and Table S4-5**). Dysregulation in *UBE2L3, IL-6, STAT4, TYK2, TNFAIP3* and *PTPN2* were found in both relapse and remission individuals, have been previously associated with JIA[21, 22]. We examined through Cytoscape and Reactome database for the gene associations involved in the 5 pathways (**Figure 5A-E**), (a) TCR activation, (b) apoptosis, (c) TNFα, (d) NF-kB and (e) MAPK signaling, and found considerable overlap between relapse and remission individuals. This overlap of pathways in relapse/remission JIA patients may arise from their shared susceptibility to clinical control with continued anti-TNFα therapy. Indeed, we observe similar disease centric pathway persistence in a separate cohort of JIA patient (n=4) that are responsive to anti-TNFα therapy, from the point of pre (active) till post-treatment (recent clinical inactivity) (**online supplementary Figure S6-7 and Table S3 and 6**). Despite this overlap, we detected selective divergence within these pathways (**Figure 5A-E**), with remission individuals expressing higher levels of *FYN, TNFRSF9, CASP1, TRAF1* and *IKBKE* which are involved in termination or resolution of these pathways[23–32].

**Figure 5.**
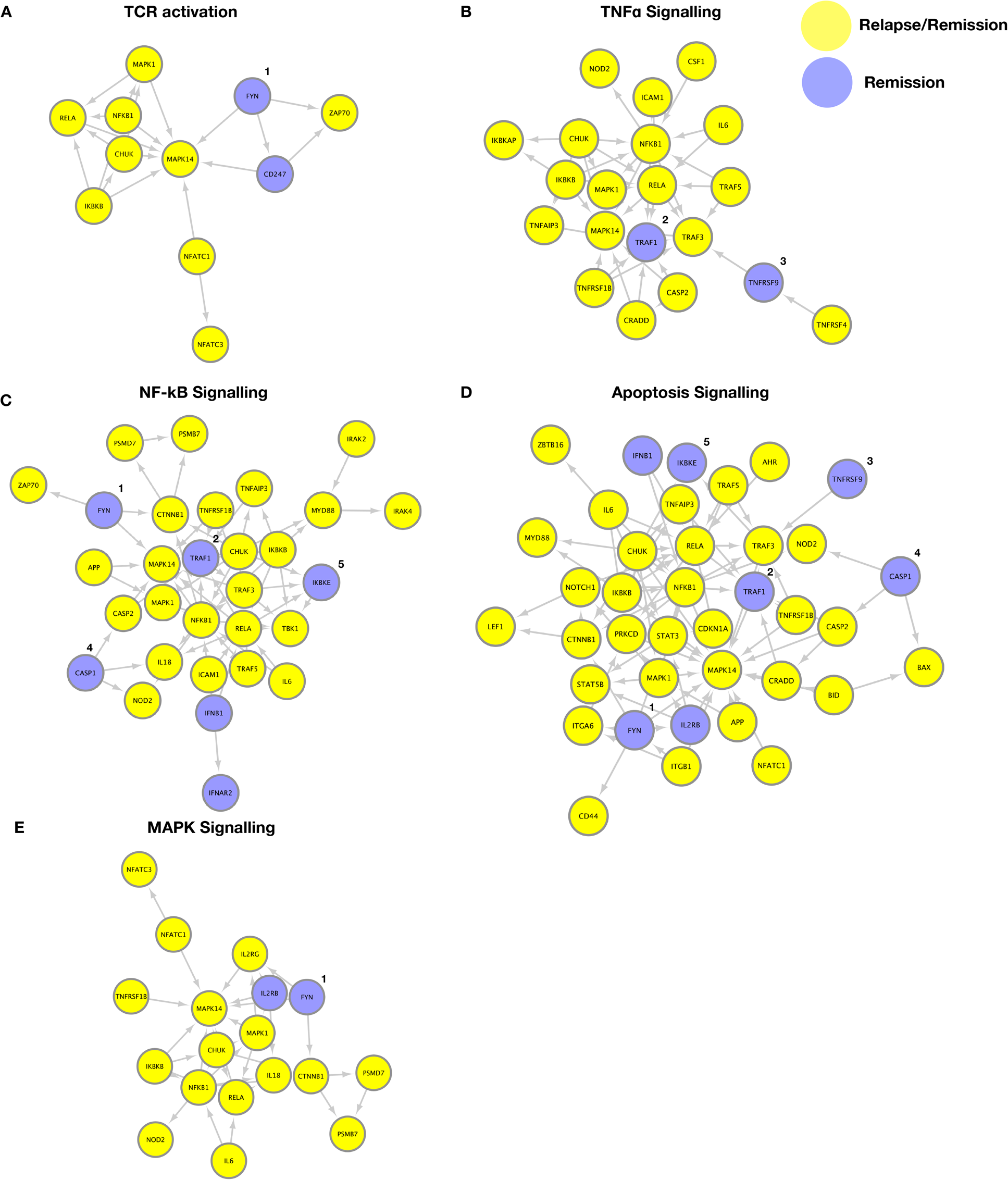
Transcriptomic divergence in disease centric pathways that persist despite therapy. Genes enriched (p <0.05, fold difference > 1.5) in patients with JIA (relapse or remission) that were persistent from prior to after therapy withdrawal as compared with healthy individuals, were exported to DAVID for functional gene-set enrichment and gene associations were constructed with Cytoscape using the Reactome database. Five major pathways were dysregulated in relapse and remission patients with JIA compared to healthy controls: (**A**) TCR activation, (**B**) TNFα signaling, (**C**) NF-kB signaling, (**D**) apoptosis, (**E**) MAPK signaling (yellow=relapse/remission, blue= remission only). Genes enriched in remission individuals include *^1^ Fyn, ^2^TRAF1, ^3^TNFRSF9, ^4^CASP1* and *^5^IKBKE*.

### CD4^+^CD45RA^−^TNFα^+^ discriminates clinical fate prior to withdrawal of therapy

We explored whether disease or remission duration of JIA patients (n=39) prior to study enrolment, could differentiate clinical fate, and found no significant difference between relapse or remission JIA patients (**Figure 6A-D**). As we have shown that an inflammatory memory subset of CD4^+^CD45RA^−^TNFα^+^ is present in relapse individuals prior to therapy withdrawal, we tested whether this could afford for discrimination in clinical fate. There was a significant difference (p < 0.001) in the ratio of CD45RA^−^TNFα^+^/CD45RA^+^TNFα^+^ cells in relapse as compared to remission individuals prior to therapy withdrawal, allowing for a ROC curve of AUC=0.9394 (**Figure 6E-F**). As age could be a serious potential cofounder for immunological memory, we determined that in the relevant age groups (7-14 yrs), there was no significant difference in CD45RA^−^ or CD45RA^−^TNFα^+^ cells among healthy individuals (n=56) (**Figure 6G-H**). Relapse individuals had significantly higher CD45RA^−^TNFα^+^ cells as compared across all the relevant age groups in healthy individuals (**Figure 6H**).

**Figure 6.**
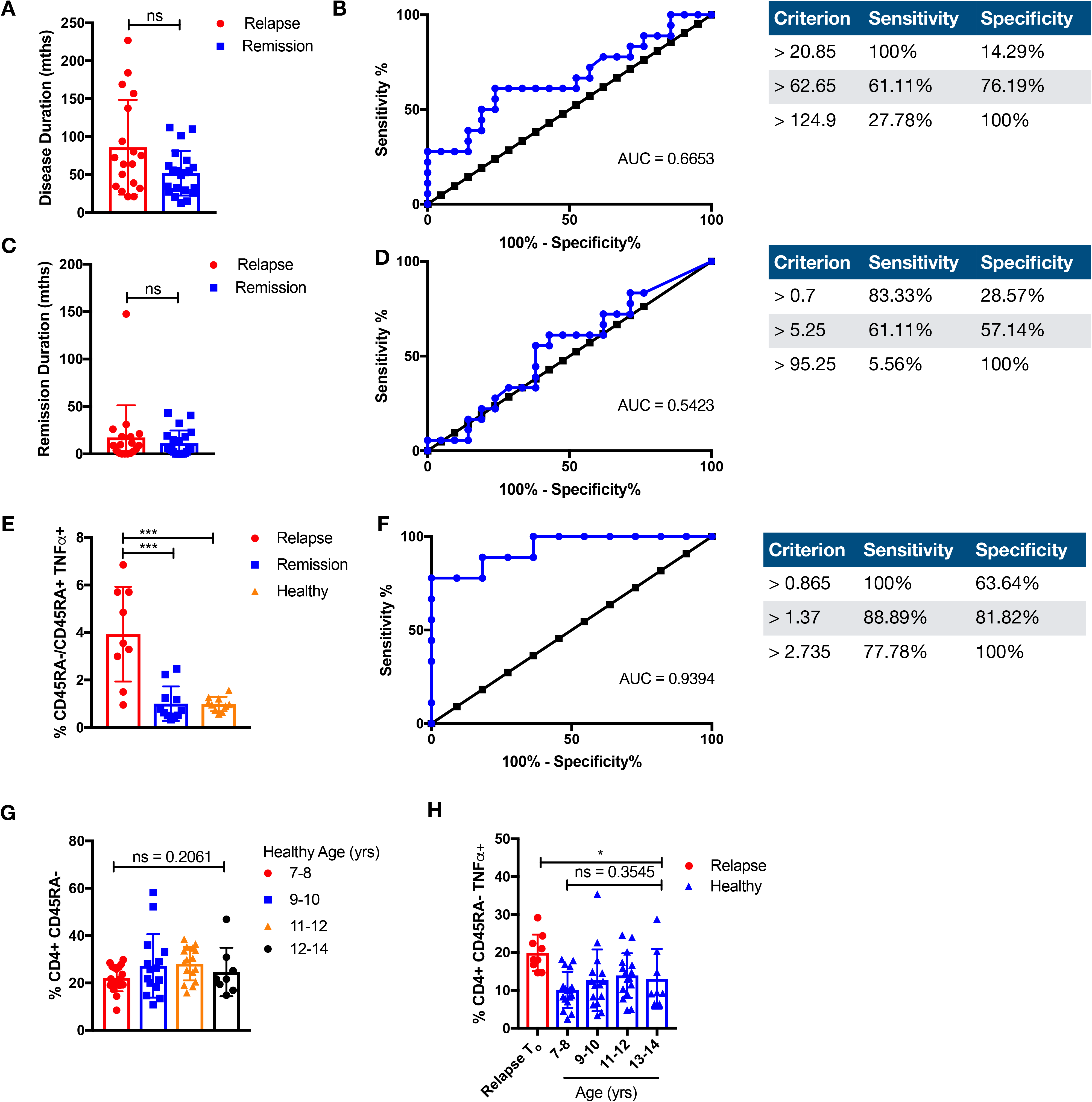
CD4^+^CD45RA^−^TNFα^+^ discriminates clinical fate. Duration and ROC of (**A-B**) disease (months) or (**C-D**) remission (months) prior to study enrolment is compared among relapse (n=18) or remission (n=21) individuals. (**E**) The ratio of memory CD4^+^CD45RA^−^ TNFα^+^ over naïve CD4^+^CD45RA^+^TNFα^+^ of relapse or remission prior to withdrawal and healthy individuals is compared. Comparison of cellular subsets performed with Mann-Whitney U, two tail test, means ± S.D., * p < 0.05. (**F**) ROC of ratio of CD4^+^CD45RA^−^ TNFα^+^ over CD4^+^CD45RA^+^TNFα^+^ of relapse or remission prior to withdrawal. (**G**) CD3^+^CD4^+^CD45RA^−^CD45RO^+^ T cells were compared in healthy paediatric controls (n=56) across the relevant age groups 7-8yrs (n=17), 9-10yrs (n=15), 11-12yrs (n=16) and 12-14yrs (n=8) or (**H**) against relapse (n=9) prior to therapy withdrawal for CD3^+^CD4^+^CD45RA^−^ TNFα^+^. Comparison of cellular subsets performed with Kruskal-Wallis test, means ± S.D.

## CONCLUSIONS

With a large proportion of JIA patients achieving clinical inactivity as a result of efficacious treatment with anti-TNFα biologics[2], it becomes increasing pertinent to address the lack of definitive withdrawal guidelines. Here, we investigated with CyToF the heterogenous CD4 landscape of patients who achieved clinical inactivity prior to therapy withdrawal. We have identified for the first time an inflammatory CD4 memory subset (CD3^+^CD4^+^CD45RA^−^ TNFα^+^) that remains elevated in JIA patients prior to relapse, and could notably discriminate clinical fate prior to therapy withdrawal (AUC=0.939).

Remarkably the presence of this inflammatory subset despite therapy, was associated with a deficit in immune checkpoint (PD1^−^CD152^−^). This is consistent with the phenomenon of irAEs (rheumatic immune-related adverse events), where the application of immune checkpoint therapy (anti-PD1/anti-CD152) in cancer results in rheumatic diseases[33]. The presence of this inflammatory subset was further verified when we separately compared relapse against healthy non-JIA individuals. The CD4^+^TNFα^+^ healthy landscape helped reveal the subclinical diversification of T-effector mechanisms in relapse individuals, with the emergence of CD3^+^CD4^+^CD45RA^−^TNFα^+^PD1^−^CD152^−^ T cells that are IL-6^+^. Particularly, CD4^+^CD45RA^−^ T cells that are IL-6^+^ express higher levels of TNFα. Recently, a case series of 3 patients with cancer and developed severe polyarthritis following immune blockade therapy, reported successful treatment with tocilizumab (anti-IL-6)[34]. This reflects a level of commonality between inflammatory and disease resolution mechanisms operating in autoimmune disorders and cancer.

We further examined a separate cohort of JIA patients that developed flare or remained in remission 8 months into withdrawal of therapy, and verified the overt presence of T-effector diversification (CD45RA^−^TNFα^+^IL-6^+^) during flare. Whereas in patients that continue to remain in remission 8 months into withdrawal, there was no differential display of any CD45RA^−^TNFα^+^ subsets as compared to the healthy CD4 landscape, though a CD4^+^CXCR3^+^CCR6^+^ subset was detected. It remains speculative if the CD4^+^CXCR3^+^CCR6^+^ subset seen in remission patients is an early disease driven subset or side-effect of therapy as it extends beyond the scope of the study, and will require longer follow-up duration. In rheumatoid arthritis, the CD4^+^CXCR3^+^CCR6^+^ subset is known express high levels of IFNγ with poor secretion of IL-17A[35], though we did not observe any associated cytokine profile in our remission patients.

In patients prior to relapse, the prevalence of the inflammatory CD4^+^CD45RA^−^TNFα^+^ T cells within the T-effector compartment is parallel by a corresponding increase in CD4^+^CD45RA^−^CD152^+^ Tregs in the regulatory arm. This suggests there is a compensatory regulatory response towards subclinical inflammation prior to relapse, though it remains to be seen whether the suppression suffices or is defective. Others have shown that synovium T-effectors are resistant to Treg suppression[19], which can be alleviated in JIA patients undergoing anti-TNFα therapy[36]. Though it is noted that in a subset of patients, in-vitro blockade of IL-6^+^ additionally alleviated T-effector resistance to Treg suppression[36].

Transcriptomic profiling of immunological genes in response to TCR signaling reveals 5 common disease centric pathways (TCR activation, apoptosis, TNFα, NF-kB and MAPK signaling) in relapse/remission patients as compared to healthy controls. Separately the same dysregulated pathways were also detected in JIA patients that are treatment naïve and later responsive to anti-TNFα therapy. This suggests that the disease pathways affected are likely an indication of susceptibility to anti-TNFα treatment. As the afflicted pathways remains dysregulated prior and after therapy withdrawal in both relapse and remission patients, exogenous therapy by sequestration of TNFα may accomplish little to reestablish healthy physiological TCR response. Despite this, we detected selective divergence in certain genes (e.g. *TRAF1*) with these pathways that may aid in termination or resolution of TCR induced signaling for remission individuals. Notably, disease association within the *TRAF1*-C5 locus[37–39], and epigenetic dysregulation within the *TRAF1* locus of CD4 T cells have been detected in JIA patients[40]. Studies using *TRAF1^−/−^* mice revealed TRAF1 negative regulatory role in T cells response to TCR and TNFα signaling[24]. *TRAF1^−/−^* T cells show enhanced proliferation in response to TCR and TNFα stimulation resulting in NF-kB and AP-1 activation[24].

In summary, by applying a combination of high dimensionality technologies, we have identified functional perturbations of the immunome in patients with arthritis who will relapse upon withdrawal for anti-TNFα therapy. These aberrations show that relapse of clinical disease relies on a foundation of complex and diverse interlacing immune mechanisms, which affect both the effector and regular arms of adaptive T cell immunity. Our findings have an immediate translational valency. Indeed, they are clinically relevant as a contribution to precision medicine [41] by offering a tool to affect clinical management. They are mechanistically relevant as, altogether, define a cluster of deranged mechanisms which can be target of focused intervention.

## Supporting information

Supplementary Test

## Contributors

JY Leong performed the CyToF, sorting and Nanostring experiments and analysis. P Chen and SL Poh helped with the CyToF and Nanostring runs. F Ally, C Chua, SN Hazirah helped with mRNA processing and Nanostring. L Pan helped write the R scripts for the CyToF analysis. L Lai helped with the CyToF. LT Bathi helped with sample processing. D Lovell, JG Yeo, ESC Lee, T Arkachaisri and PRSCG, helped with patient recruitment. S Albani was the lead principal investigator.

## Ethics approval

All samples were collected upon informed consent. All experiments were conducted according to the principles expressed in the Declaration of Helsinki, and IRB approved by the Sanford-Burnham or KK Women’s and Children’s Hospital IRB Committee.

