## Supplementary Figure for "An immunome perturbation is present in juvenile idiopathic arthritis patients who are in remission and will relapse upon anti-TNFα withdrawal"

**Supplementary Figure S1**

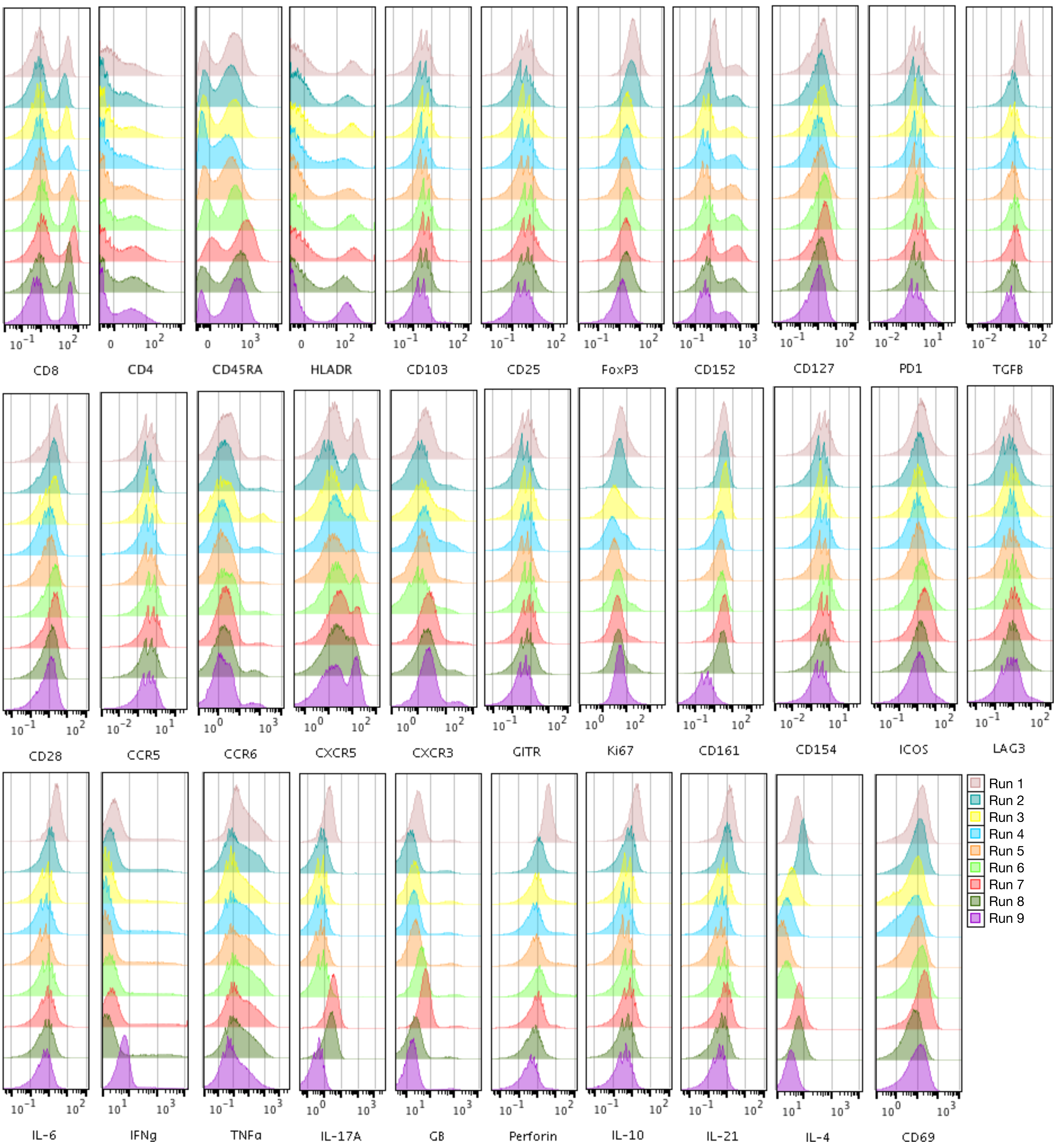

**Supplementary Figure S2**

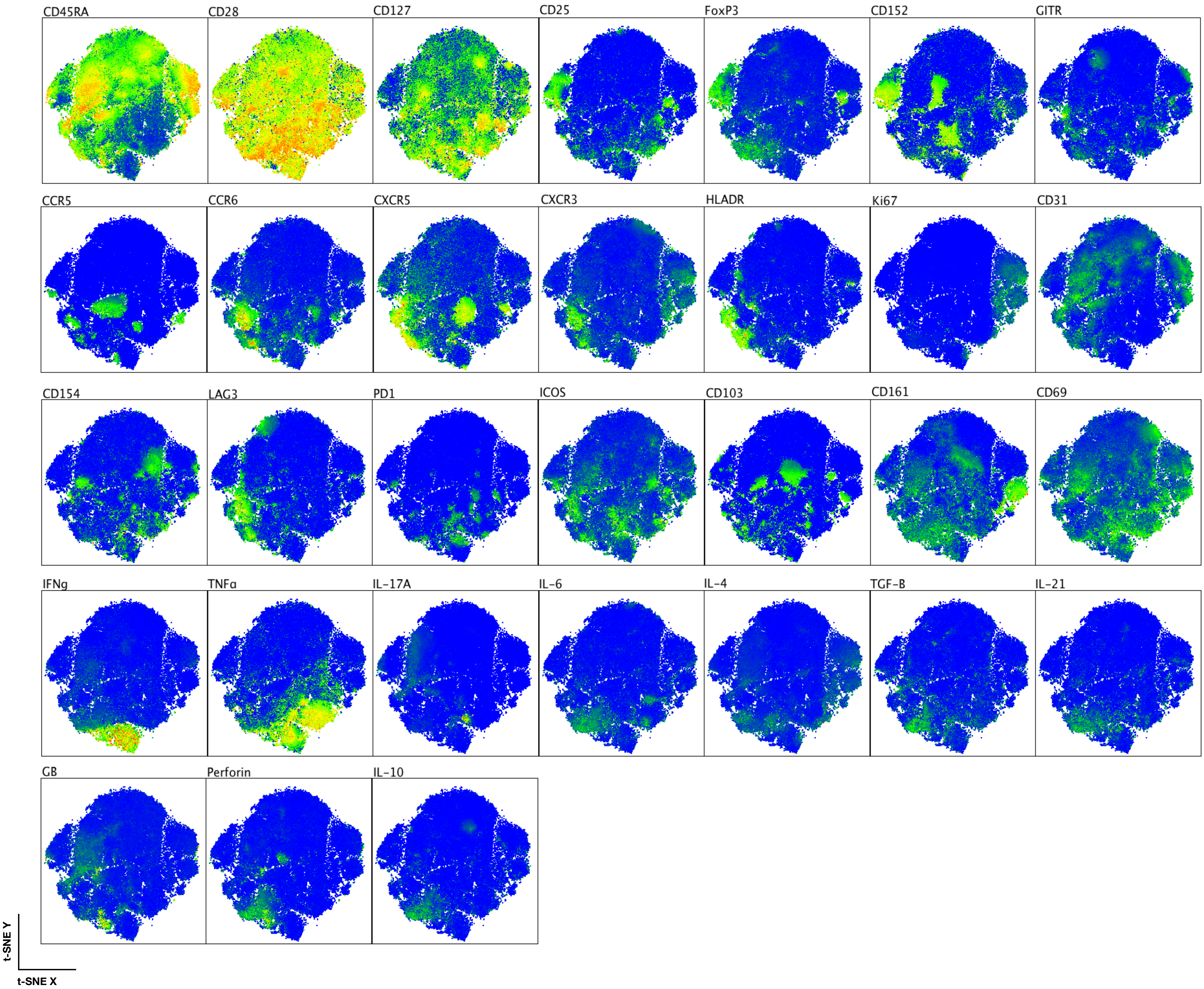

### Supplementary Figure S3

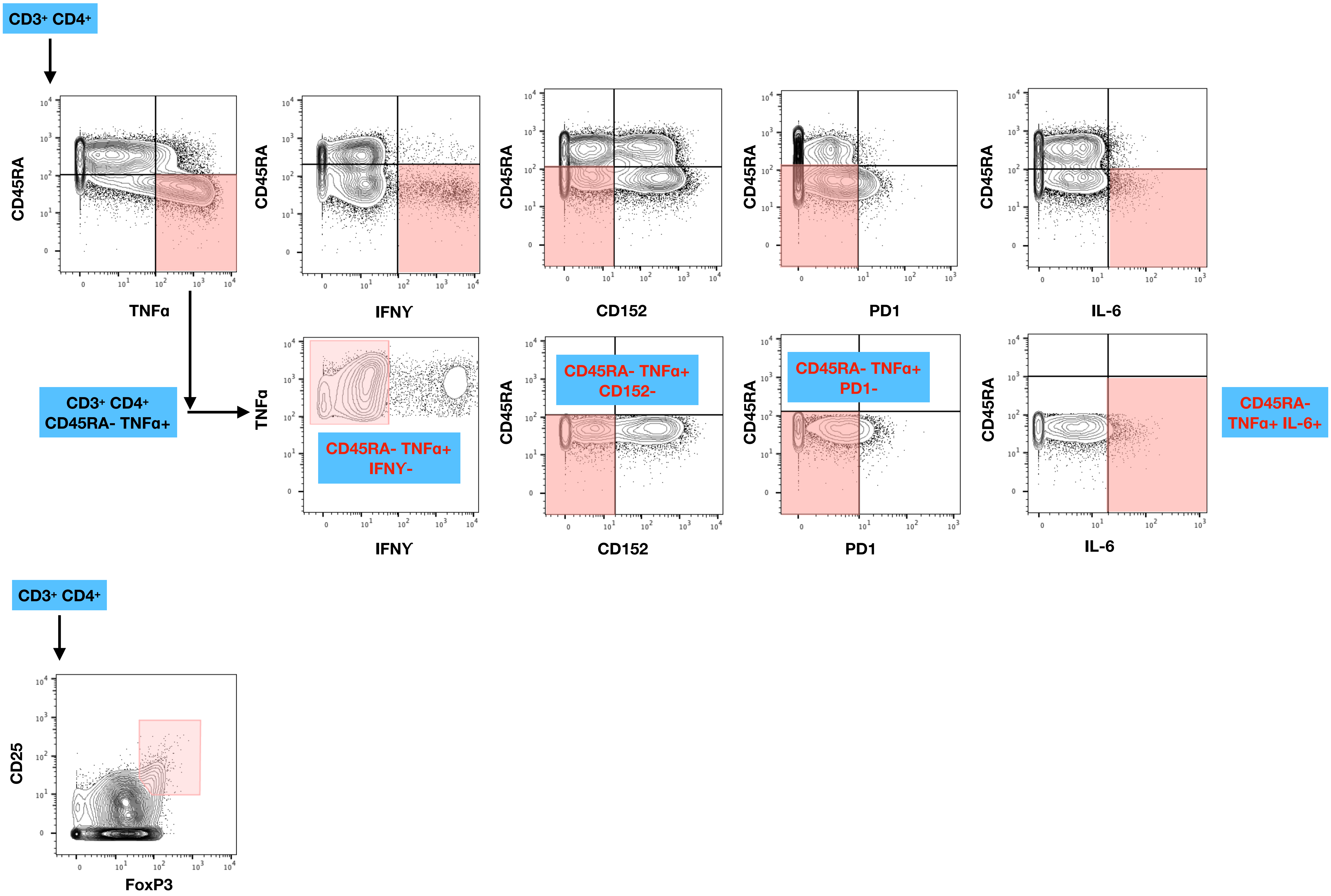

### Supplementary Figure S4

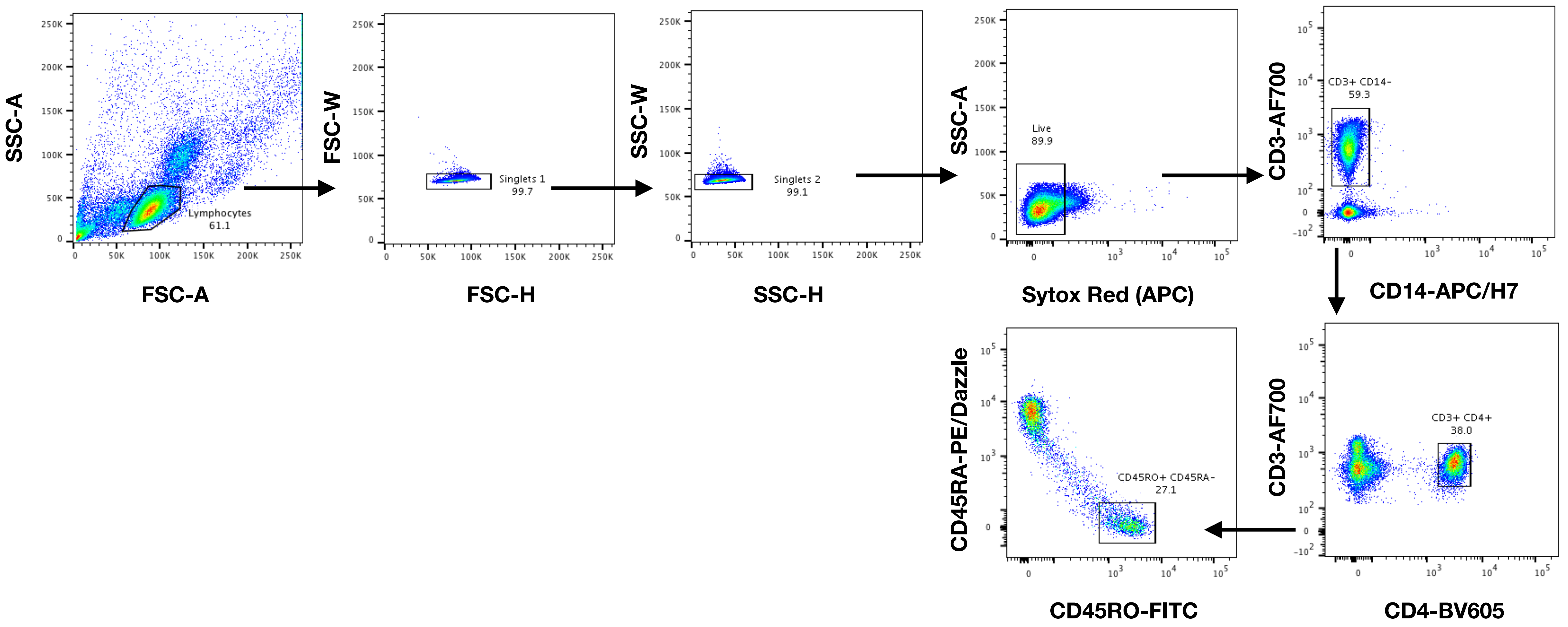

#### Supplementary Figure S5

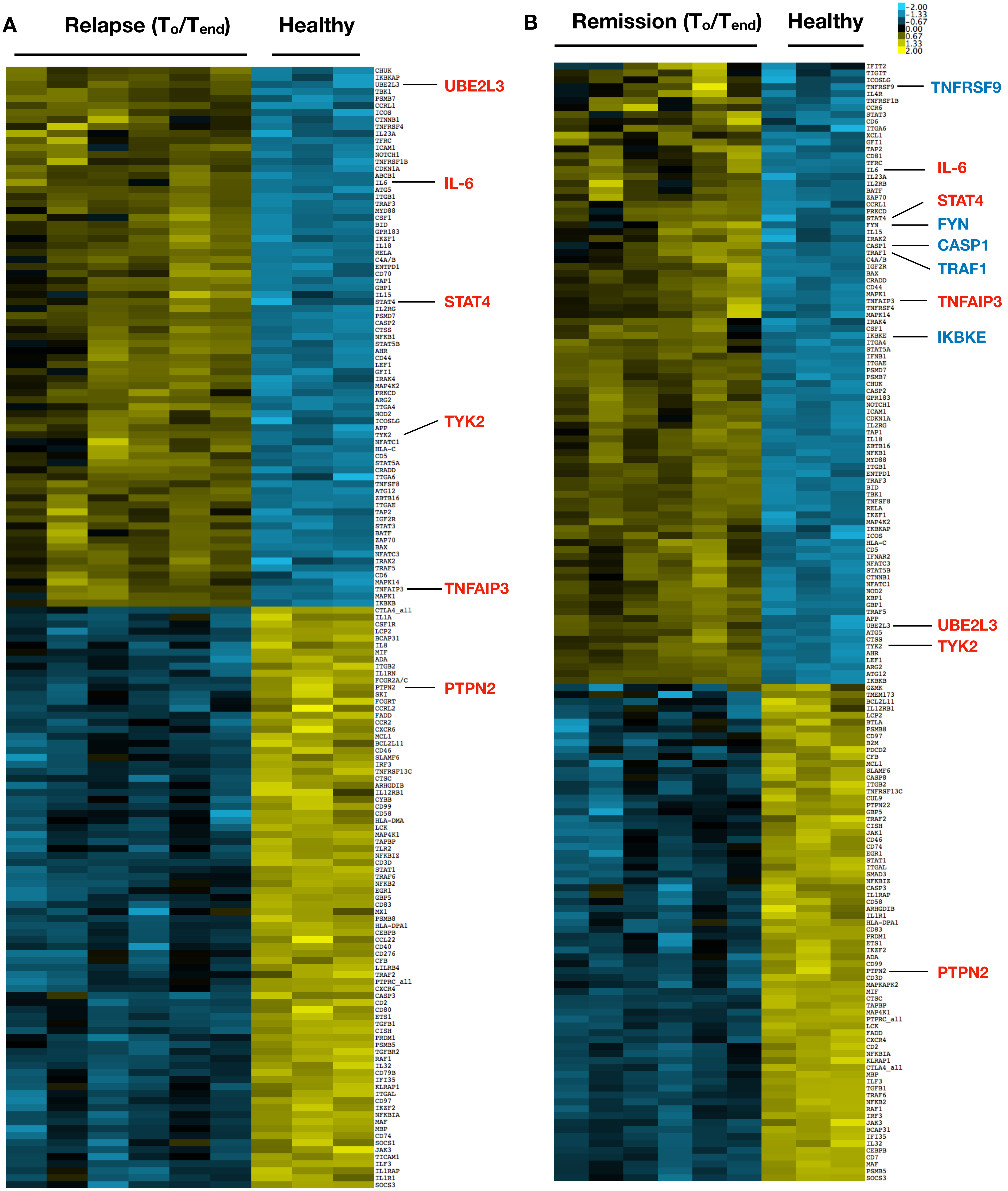

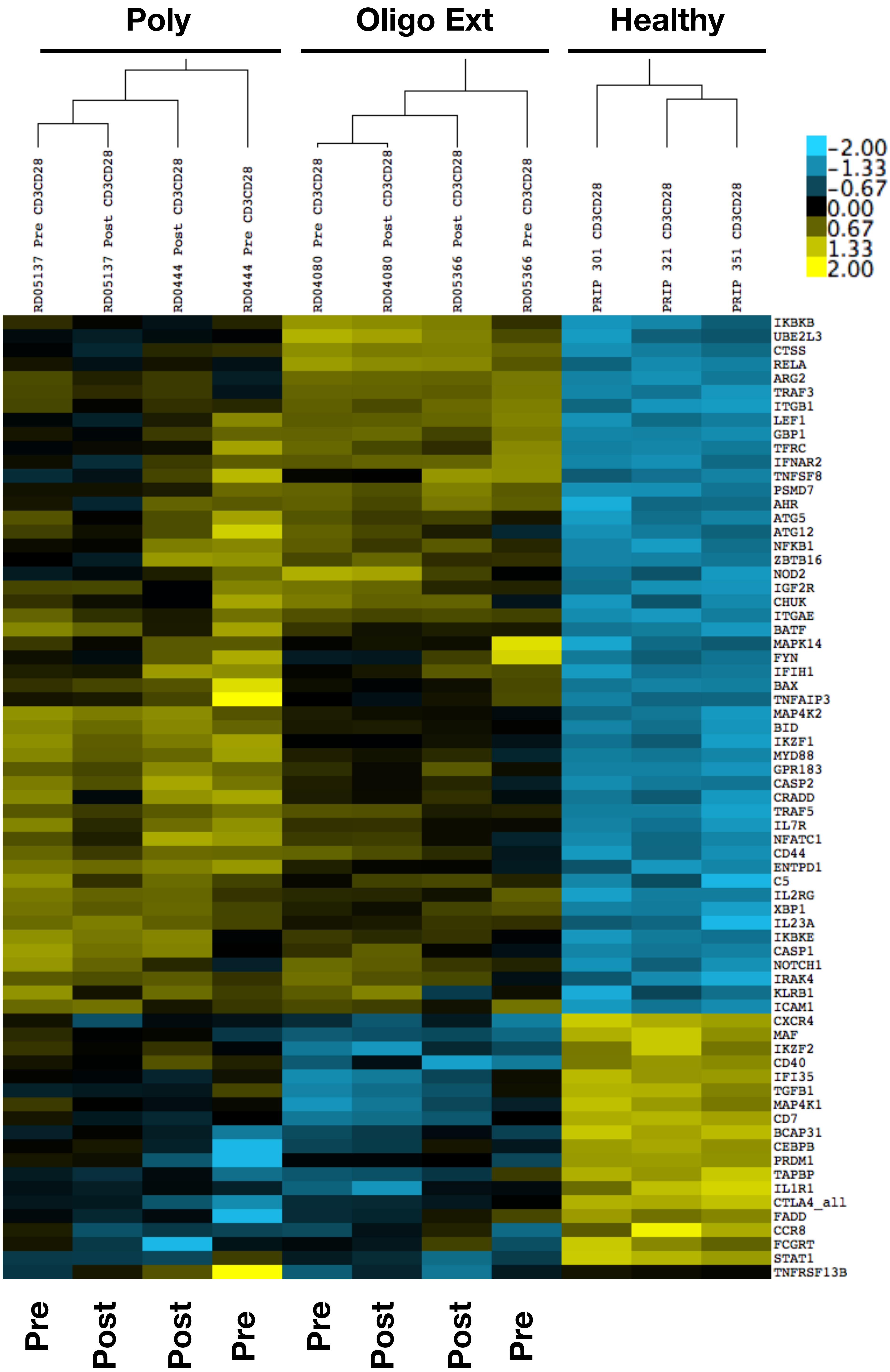
Poly
Oligo Ext
Healthy
RD05137 Pre CD3CD28
RD05137 Post CD3CD28
RD0444 Post CD3CD28
RD0444 Pre CD3CD28
RD04080 Pre CD3CD28
RD04080 Post CD3CD28
RD05366 Post CD3CD28
RD05366 Pre CD3CD28
PRIP 301 CD3CD28
PRIP 321 CD3CD28
PRIP 351 CD3CD28
-2.00
-1.33
-0.67
0.00
0.67
1.33
2.00
IKBKB
UBE2L3
CTSS
RELA
ARG2
TRAF3
ITGB1
LEF1
GBP1
TFRC
IFNAR2
TNFSF8
PSMD7
AHR
ATG5
ATG12
NFKB1
ZBTB16
NOD2
IGF2R
CHUK
ITGAE
BATF
MAPK14
FYN
IFIH1
BAX
TNFAIP3
MAP4K2
BID
IKZF1
MYD88
GPR183
CASP2
CRADD
TRAF5
IL7R
NFATC1
CD44
ENTPD1
C5
IL2RG
XBP1
IL23A
IKBKE
CASP1
NOTCH1
IRAK4
KLRB1
ICAM1
CXCR4
MAF
IKZF2
CD40
IFI35
TGFB1
MAP4K1
CD7
BCAP31
CEBPB
PRDM1
TAPBP
IL1R1
CTLA4_all
FADD
CCR8
FCGRT
STAT1
TNFRSF13B
Pre
Post
Post
Pre
Pre
Post
Post
Pre

#### Supplementary Figure S7

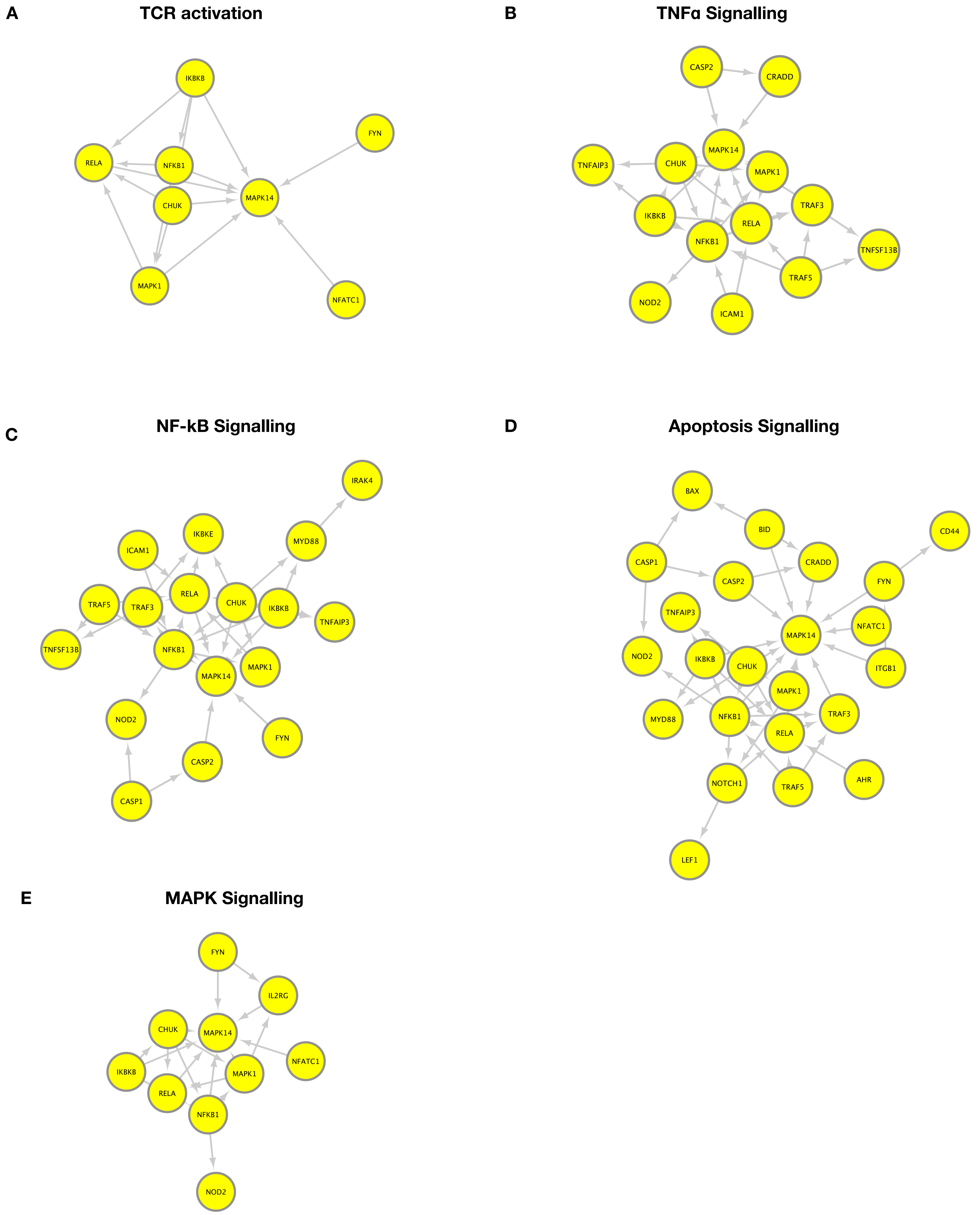

#### Supplementary Figure S8

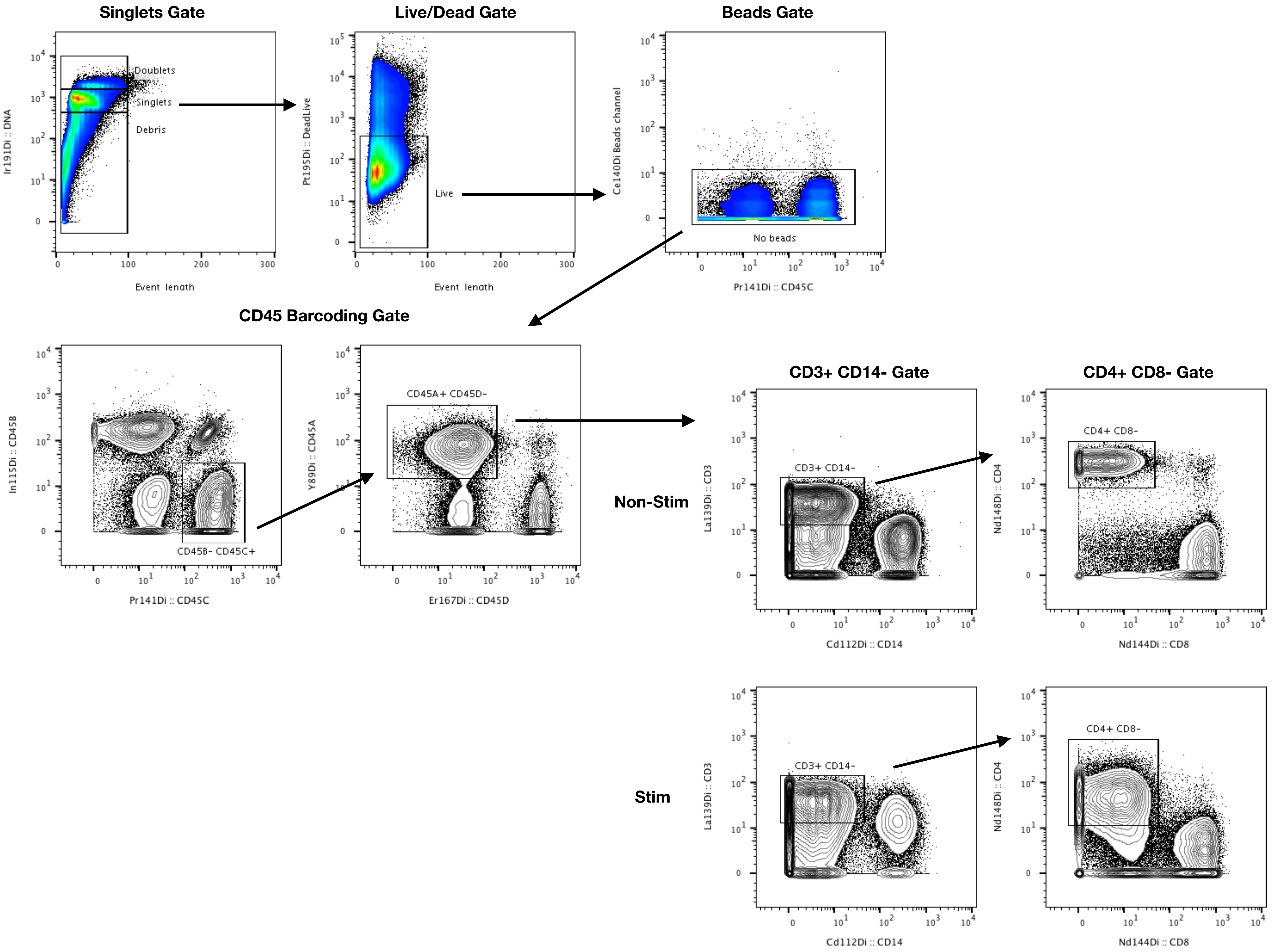
