## Supplementary Table for "An immunome perturbation is present in juvenile idiopathic arthritis patients who are in remission and will relapse upon anti-TNFα withdrawal"

### Supplementary Table S1

| No. | Individuals | JIA Subtype | Gender | Medication History | CyToF | Nanostring |
|---|---|---|---|---|---|---|
| 1 | Relapse 1011 ($T_o$) | Poly RF - | F | etanercept, no MTX | ✓ | |
| 2 | Relapse 1044 ($T_o$) | Poly RF - | F | etanercept, no MTX | ✓ | |
| 3 | Relapse 2020 ($T_o$) | Poly RF - | F | adalimumab, no MTX | ✓ | |
| 4 | Relapse 2042 ($T_o$) | Poly RF - | F | etanercept, no MTX | ✓ | |
| 5 | Relapse 4090 ($T_o$) | Poly RF - | F | etanercept, no MTX | ✓ | |
| 6 | Relapse 5031 ($T_o$) | Poly RF - | F | adalimumab, yes MTX | ✓ | |
| 7 | Relapse 6129 ($T_o$) | Poly RF - | F | adalimumab, no MTX | ✓ | |
| 8 | Relapse 7063 ($T_o$) | Poly RF - | F | etanercept, yes MTX | ✓ | |
| 9 | Relapse 8120 ($T_o$) | Poly RF - | F | etanercept, no MTX | ✓ | |
| 10 | Relapse 5137 ($T_o$) | Poly RF - | M | infliximab, yes MTX | | ✓ |
| 11 | Relapse 10107 ($T_o$) | Poly RF - | F | etanercept, no MTX | | ✓ |
| 12 | Relapse 1006 ($T_o$) | Poly RF - | F | etanercept, no MTX | | ✓ |
| 1 | Relapse 1001 ($T_{end}$) | Poly RF - | F | etanercept, no MTX | ✓ | |
| 2 | Relapse 1003 ($T_{end}$) | Poly RF - | F | etanercept, yes MTX | ✓ | |
| 3 | Relapse 1006 ($T_{end}$) | Poly RF - | F | etanercept, no MTX | ✓ | |
| 4 | Relapse 1011 ($T_{end}$) | Poly RF - | F | etanercept, no MTX | ✓ | |
| 5 | Relapse 1044 ($T_{end}$) | Poly RF - | F | etanercept, no MTX | ✓ | |
| 6 | Relapse 3066 ($T_{end}$) | Poly RF - | F | etanercept, no MTX | ✓ | |
| 7 | Relapse 6049 ($T_{end}$) | Poly RF - | F | etanercept, no MTX | ✓ | ✓ |
| 8 | Relapse 7063 ($T_{end}$) | Poly RF - | F | etanercept, yes MTX | ✓ | |
| 9 | Relapse 10107 ($T_{end}$) | Poly RF - | F | etanercept, no MTX | ✓ | |
| 10 | Relapse 3062 ($T_{end}$) | Poly RF - | M | infliximab, yes MTX | | ✓ |
| 11 | Relapse 6048 ($T_{end}$) | Poly RF - | F | etanercept, yes MTX | | ✓ |

| No. | Individuals | JIA Subtype | Gender | Medication History | CyToF | Nanostring |
|---|---|---|---|---|---|---|
| 1 | Remission 1018 ($T_o$) | Poly RF - | F | etanercept, no MTX | ✓ | |
| 2 | Remission 1034 ($T_o$) | Poly RF - | M | etanercept, no MTX | ✓ | |
| 3 | Remission 1124 ($T_o$) | Poly RF - | F | etanercept, no MTX | ✓ | |
| 4 | Remission 2022 ($T_o$) | Poly RF + | F | etanercept, yes MTX | ✓ | ✓ |
| 5 | Remission 2114 ($T_o$) | Poly RF - | F | etanercept, no MTX | ✓ | |
| 6 | Remission 4014 ($T_o$) | Poly RF - | F | etanercept, no MTX | ✓ | |
| 7 | Remission 5121 ($T_o$) | Poly RF - | M | etanercept, yes MTX | ✓ | |
| 8 | Remission 6128 ($T_o$) | Poly RF - | M | etanercept, yes MTX | ✓ | |
| 9 | Remission 7056 ($T_o$) | Poly RF - | F | etanercept, no MTX | ✓ | |
| 10 | Remission 8130 ($T_o$) | Poly RF - | F | adalimumab, yes MTX | ✓ | |
| 11 | Remission 15122 ($T_o$) | Poly RF + | F | etanercept, no MTX | ✓ | |
| 12 | Remission 5127 ($T_o$) | Poly RF - | F | etanercept, yes MTX | | ✓ |
| 13 | Remission 4073 ($T_o$) | Poly RF + | F | adalimumab, no MTX | | ✓ |
| 1 | Remission 2022 ($T_{end}$) | Poly RF + | F | etanercept, yes MTX | ✓ | |
| 2 | Remission 5026 ($T_{end}$) | Poly RF - | M | etanercept, yes MTX | ✓ | |
| 3 | Remission 8054 ($T_{end}$) | Poly RF - | F | etanercept, no MTX | ✓ | |
| 4 | Remission 8131 ($T_{end}$) | Poly RF - | M | etanercept, no MTX | ✓ | |
| 5 | Remission 9086 ($T_{end}$) | Poly RF - | F | etanercept, no MTX | ✓ | ✓ |
| 6 | Remission 9098 ($T_{end}$) | Poly RF - | F | etanercept, no MTX | ✓ | |
| 7 | Remission 16111 ($T_{end}$) | Poly RF + | F | etanercept, yes MTX | ✓ | |
| 8 | Remission 10081 ($T_{end}$) | Poly RF + | M | etanercept, yes MTX | | ✓ |
| 9 | Remission 3103 ($T_{end}$) | Poly RF + | F | adalimumab, no MTX | | ✓ |

| | | Relapse | | Remission | | Healthy |
|---|---|---|---|---|---|---|
| | | $T_o$ (n=12) | $T_{end}$ (n=11) | $T_o$ (n=13) | $T_{end}$ (n=9) | n= 69 |
| Anti-TNFα Biologics | etanercept (%) | 8 (66.7%) | 10 (90.9%) | 11 (84.6%) | 9 (100%) | N.A. |
| | adalimumab (%) | 3 (25%) | 0 (0%) | 2 (15.4%) | 0 (0%) | N.A. |
| | Infliximab (%) | 1 (8.33%) | 1 (9.1%) | 0 (0%) | 0 (0%) | N.A. |
| Methotrexate (concurrent) (%) | | 3 (25%) | 4 (36.4%) | 5 (38.5%) | 4 (44.4%) | N.A. |
| Rheumatoid Factor+ | | 0 (0%) | 0 | 3 (23.1%) | 4 (44.4%) | N.A. |
| Average age (yrs) ± Std Dev | | 14.2 ± 4.2 | 12.5 ± 5.1 | 9.0 ± 3.4 | 12.6 ± 4.2 | 9 ± 3 |
| Female (n): Male (n) | | 11:1 | 10:1 | 10:3 | 6:3 | 10:59 |

### Supplementary Table S2

| Patient ID | Date | Disease | age | Disease Duration (yrs) | Gender | Joints | ESR | CRP | Medication history |
|---|---|---|---|---|---|---|---|---|---|
| RD00444 Pre | 25/7/16 | Poly JIA, RF+ | 16.5 | 11.8 | F | 5 | Not done | Not done | Sulfasalazine, MTX, Folic acid, HCQ |
| RD00444 Post | 10/2/17 | Poly JIA, RF+ | 17.0 | 12.4 | F | 0 | 48 | 2 | MTX, Folic acid, Enbrel |
| RD05137 Pre | 11/3/16 | Poly JIA, RF+ | 10.9 | 0.1 | F | 4 | 26 | 1.5 | - |
| RD05137 Post | 15/9/16 | Poly JIA, RF+ | 11.4 | 0.6 | F | 0 | 7 | <0.2 | Enbrel, MTX, Folic acid |
| RD04080 Pre | 13/4/16 | Oligo JIA, Extended | 13.6 | 2.4 | F | 4 | 47 | 18.2 | MTX, Prednisolone, Omeprazole, Folic acid |
| RD04080 Post | 26/10/16 | Oligo JIA, Extended | 14.1 | 2.9 | F | 0 | 21 | 5.7 | MTX, Folic acid, Enbrel |
| RD05336 Pre | 4/8/16 | Oligo JIA, Extended | 1.9 | 0.1 | F | 8 | 97 | 49.3 | Brufen |
| RD05336 Post | 10/2/17 | Oligo JIA, Extended | 2.4 | 0.6 | F | 0 | 8 | 0.4 | Enbrel |

### Supplementary Table S3

| Targets | Metal Channel | Staining | Clone | Antibody Vendor/Catalogue number |
|---|---|---|---|---|
| Lineage markers | | | | |
| CD3 | 139 | Surface/Intra | UCHT1 | Biolegend (300402) |
| CD4 | 148 | Surface/Intra | SK3 | Biolegend (344625) |
| CD8 | 144 | Surface | SK1 | Biolegend (344727) |
| CD11b | 161 | Surface | ICRF44 | Biolegend (301302) |
| CD16 | 209 | Surface | 3G8 | Fluidigm (3209002B) |
| CD14 | 112/114 | Surface | M5E2 | Biolegend (301843) |
| T helper subsets | | | | |
| IL-4 | 156 | Intra | 8D4-8 | Biolegend (500707) |
| IFN-g | 168 | Intra | B27 | Biolegend (506513) |
| IL-17A | 169 | Intra | BL168 | Biolegend (512302) |
| IL-21 | 151 | Intra | 3A4-N2 | Biolegend (513009) |
| CD161 | 157 | Surface | HP-3G10 | Biolegend (339902) |
| T cell functional | | | | |
| CD45RA | 171 | Surface | HI100 | Biolegend (304102) |
| CD69 | 176 | Surface | FN50 | Biolegend (310902) |
| CD28 | 146 | Surface | CD28.2 | Biolegend (302923) |
| CD152 (CTLA4) | 155 | Intra | BNI3 | Biolegend (555851) |
| CD154 (CD40L) | 149 | Surface | 24-31 | Biolegend (310835) |
| HLA-DR | 143 | Surface | L243 | Biolegend (307612) |
| LAG3 | 159 | Surface | 17B4 | Abcam (ab40466) |
| PD1 | 147 | Surface | EH12.2H7 | Biolegend (329941) |
| Ki67 | 166 | Intra | 20Raj1 | Thermofisher/ebioscience (14-5699-82) |
| ICOS | 154 | Surface | C398.4A | Biolegend (313512) |
| CD31 | 172 | Surface | WM59 | Biolegend (303102) |
| CD103 | 142 | Surface | B-Ly7 | Thermofisher/ebioscience (14-1038-82) |
| Chemokine receptors | | | | |
| CXCR3 | 163 | Surface | G025H7 | Biolegend (353718) |
| CXCR5 | 160 | Surface | RF8B2 | BD biosciences (552032) |
| CCR5 | 145 | Surface | NP-6G4 | Abcam (ab115738) |
| CCR6 | 170 | Surface | G034E3 | Biolegend (353402) |
| Treg markers | | | | |
| CD25 | 150 | Surface | M-A251 | BD biosciences (555429) |
| CD127 | 153 | Surface | A019D5 | Biolegend (351302) |
| FoxP3 | 165 | Intra | PCH10L | Thermofisher/ebioscience (14-4776-82) |
| GITR | 164 | Surface | 621 | Biolegend (311602) |
| TGF-B (LAP) | 175 | Surface/Intra | TW4-2F8 | Biolegend (349602) |
| IL-10 | 158 | Intra | JES3-9D7 | Biolegend (501402) |
| Cytokines/Enzymes | | | | |
| TNFα | 152 | Intra | Mab11 | Biolegend (502902) |
| IL-6 | 162 | Intra | MQ2-13A5 | Thermofisher/ebioscience (16-7069-85) |
| Granzyme B | 173 | Intra | CLB-GB11 | Abcam (ab103159) |
| Perforin | 174 | Intra | B-D48 | Abcam (ab47225) |
| Barcodes | | | | |
| CD45-A | 89 | Surface | HI30 | Fluidigm (3089003B) |
| CD45-B,C or D | 115, 141, 167 | Surface | HI30 | Biolegend (304002) |
| Live/Dead /Singlets | | | | |
| DNA (Singlets) | 191/193 | - | Nil | Fluidigm Cell-ID Intercalator-Ir (201192B) |
| Cisplatin (Live/Dead) | 195 | - | Nil | Sigma-aldrich (479306-1G) |

### Supplementary Table S4

| Category | Term | Count | % | P-Value | Benjamin |
|---|---|---|---|---|---|
| **TCR activation** | | | | | |
| KEGG_PATHWAY | T cell receptor signaling pathway | 10 | 13.2 | 4.7E-07 | 3.9E-06 |
| GOTERM_BP_DIRECT | T cell receptor signaling pathway | 8 | 10.5 | 4.3E-06 | 3.5E-04 |
| REACTOME_PATHWAY | R-HSA-202424 (Downstream TCR signaling) | 6 | 7.9 | 5.6E-04 | 9.4E-03 |
| BIOCARTA | T Cell Receptor Signaling Pathway | 5 | 6.6 | 4.6E-02 | 2.4E-01 |
| **Apoptosis** | | | | | |
| UP_KEYWORDS | Apoptosis | 15 | 19.7 | 7.3E-09 | 2.6E-07 |
| GOTERM_BP_DIRECT | apoptotic process | 14 | 18.4 | 1.2E-06 | 1.4E-04 |
| GOTERM_BP_DIRECT | negative regulation of apoptotic process | 11 | 14.5 | 3.3E-05 | 1.8E-03 |
| GOTERM_BP_DIRECT | positive regulation of apoptotic process | 10 | 13.2 | 7.4E-06 | 4.6E-04 |
| GOTERM_BP_DIRECT | regulation of apoptotic process | 9 | 11.8 | 4.8E-06 | 3.5E-04 |
| KEGG_PATHWAY | Apoptosis | 6 | 7.9 | 2.9E-04 | 1.2E-03 |
| GOTERM_BP_DIRECT | apoptotic signaling pathway | 5 | 6.6 | 2.8E-04 | 9.7E-03 |
| GOTERM_BP_DIRECT | activation of cysteine-type endopeptidase activity involved in apoptotic process | 4 | 5.3 | 6.1E-03 | 9.6E-02 |
| BIOCARTA | Neuropeptides VIP and PACAP inhibit the apoptosis of activated T cells | 4 | 5.3 | 4.6E-02 | 2.5E-01 |
| BIOCARTA | Induction of apoptosis through DR3 and DR4/5 Death Receptors | 4 | 5.3 | 8.2E-02 | 3.6E-01 |
| **TNF-alpha signalling** | | | | | |
| KEGG_PATHWAY | TNF signaling pathway | 15 | 19.7 | 6.7E-13 | 1.0E-10 |
| BIOCARTA | TNFR2 Signaling Pathway | 8 | 10.5 | 4.8E-07 | 7.4E-05 |
| BIOCARTA | TNF/Stress Related Signaling | 8 | 10.5 | 6.2E-06 | 3.2E-04 |
| GOTERM_BP_DIRECT | tumor necrosis factor-mediated signaling pathway | 7 | 9.2 | 1.4E-05 | 8.5E-04 |
| GOTERM_BP_DIRECT | cellular response to tumor necrosis factor | 5 | 6.6 | 1.5E-03 | 3.6E-02 |
| **NF-kB signalling** | | | | | |
| KEGG_PATHWAY | Toll-like receptor signaling pathway | 11 | 14.5 | 4.8E-08 | 5.3E-07 |
| KEGG_PATHWAY | NF-kappa B signaling pathway | 11 | 14.5 | 7.0E-09 | 1.2E-07 |
| KEGG_PATHWAY | NOD-like receptor signaling pathway | 10 | 13.2 | 1.7E-09 | 5.2E-08 |
| GOTERM_BP_DIRECT | positive regulation of NF-kappaB transcription factor activity | 10 | 13.2 | 7.5E-09 | 2.0E-06 |
| GOTERM_BP_DIRECT | positive regulation of I-kappaB kinase/NF-kappaB signaling | 9 | 11.8 | 5.9E-07 | 7.6E-05 |
| KEGG_PATHWAY | RIG-I-like receptor signaling pathway | 9 | 11.8 | 2.7E-07 | 2.7E-06 |
| REACTOME_PATHWAY | R-HSA-445989 (TAK1 activates NFkB by phosphorylation and activation of IKKs complex) | 8 | 10.5 | 7.8E-10 | 2.0E-07 |
| REACTOME_PATHWAY | R-HSA-168638 (NOD1/2 Signaling Pathway) | 7 | 9.2 | 1.2E-07 | 1.5E-05 |
| KEGG_PATHWAY | Cytosolic DNA-sensing pathway | 7 | 9.2 | 2.9E-05 | 1.6E-04 |
| BIOCARTA | NF-kB Signaling Pathway | 7 | 9.2 | 4.8E-05 | 1.5E-03 |
| REACTOME_PATHWAY | R-HSA-1810476 (RIP-mediated NFkB activation via ZBP1) | 6 | 7.9 | 2.2E-07 | 1.8E-05 |
| REACTOME_PATHWAY | R-HSA-933542 (TRAF6 mediated NF-kB activation) | 6 | 7.9 | 4.5E-07 | 2.8E-05 |
| REACTOME_PATHWAY | R-HSA-2871837 (FCERI mediated NF-kB activation) | 6 | 7.9 | 2.2E-03 | 2.7E-02 |
| GOTERM_BP_DIRECT | nucleotide-binding oligomerization domain containing signaling pathway | 5 | 6.6 | 4.3E-06 | 3.3E-04 |
| GOTERM_BP_DIRECT | I-kappaB kinase/NF-kappaB signaling | 5 | 6.6 | 1.5E-04 | 6.4E-03 |
| GOTERM_BP_DIRECT | NIK/NF-kappaB signaling | 4 | 5.3 | 3.2E-03 | 6.1E-02 |
| GOTERM_BP_DIRECT | negative regulation of NF-kappaB transcription factor activity | 4 | 5.3 | 3.9E-03 | 7.2E-02 |
| REACTOME_PATHWAY | R-HSA-5668541 (TNFR2 non-canonical NF-kB pathway) | 4 | 5.3 | 1.1E-02 | 8.9E-02 |
| **MAPK signalling** | | | | | |
| KEGG_PATHWAY | MAPK signaling pathway | 9 | 11.8 | 2.8E-03 | 8.9E-03 |
| BIOCARTA | MAPKinase Signaling Pathway | 7 | 9.2 | 5.1E-02 | 2.6E-01 |
| GOTERM_BP_DIRECT | MAPK cascade | 5 | 6.6 | 3.0E-02 | 3E-01 |
| GOTERM_BP_DIRECT | positive regulation of MAPK cascade | 4 | 5.3 | 5.7E-03 | 9.1E-02 |
| GOTERM_BP_DIRECT | activation of MAPK activity | 4 | 5.3 | 1.2E-02 | 1.6E-01 |

### Supplementary Table S5

| Category | Term | Count | % | P-Value | Benjamin |
|---|---|---|---|---|---|
| **TCR activation** | | | | | |
| KEGG_PATHWAY | T cell receptor signaling pathway | 12 | 13.3 | 1.8E-08 | 1.6E-07 |
| GOTERM_BP_DIRECT | T cell receptor signaling pathway | 10 | 11.1 | 8.8E-08 | 1.4E-05 |
| REACTOME_PATHWAY | R-HSA-202424 (Downstream TCR signaling) | 7 | 7.8 | 1.4E-04 | 3.4E-03 |
| BIOCARTA | T Cell Receptor Signaling Pathway | 7 | 7.8 | 4.6E-03 | 5.2E-02 |
| **Apoptosis** | | | | | |
| UP_KEYWORDS | Apoptosis | 18 | 20.0 | 1.2E-10 | 6.1E-09 |
| GOTERM_BP_DIRECT | apoptotic process | 17 | 18.9 | 4.2E-08 | 7.2E-06 |
| GOTERM_BP_DIRECT | regulation of apoptotic process | 12 | 13.3 | 1.6E-08 | 3.2E-06 |
| GOTERM_BP_DIRECT | negative regulation of apoptotic process | 12 | 13.3 | 2.8E-05 | 1.3E-03 |
| GOTERM_BP_DIRECT | positive regulation of apoptotic process | 11 | 12.2 | 4.2E-06 | 3.2E-04 |
| KEGG_PATHWAY | Apoptosis | 6 | 6.7 | 6.7E-04 | 2.5E-03 |
| GOTERM_BP_DIRECT | apoptotic signaling pathway | 5 | 5.6 | 5.5E-04 | 1.3E-02 |
| GOTERM_BP_DIRECT | activation of cysteine-type endopeptidase activity involved in apoptotic process | 5 | 5.6 | 9.8E-04 | 2.1E-02 |
| GOTERM_BP_DIRECT | positive regulation of apoptotic signaling pathway | 4 | 4.4 | 3.0E-04 | 9.3E-03 |
| GOTERM_BP_DIRECT | extrinsic apoptotic signaling pathway via death domain receptors | 4 | 4.4 | 1.1E-03 | 2.3E-02 |
| GOTERM_BP_DIRECT | intrinsic apoptotic signaling pathway in response to DNA damage | 4 | 4.4 | 2.0E-03 | 3.7E-02 |
| BIOCARTA | Neuropeptides VIP and PACAP inhibit the apoptosis of activated T cells | 4 | 4.4 | 6.8E-02 | 3.1E-01 |
| **TNF-alpha signalling** | | | | | |
| KEGG_PATHWAY | TNF signaling pathway | 16 | 17.8 | 5.3E-13 | 2.7E-11 |
| BIOCARTA | TNFR2 Signaling Pathway | 9 | 10.0 | 7.1E-08 | 1.1E-05 |
| BIOCARTA | TNF/Stress Related Signaling | 8 | 8.9 | 1.9E-05 | 7.6E-04 |
| GOTERM_BP_DIRECT | tumor necrosis factor-mediated signaling pathway | 7 | 7.8 | 3.8E-05 | 1.7E-03 |
| GOTERM_BP_DIRECT | cellular response to tumor necrosis factor | 6 | 6.7 | 3.0E-04 | 9.3E-03 |
| GOTERM_BP_DIRECT | regulation of tumor necrosis factor-mediated signaling pathway | 4 | 4.4 | 5.3E-04 | 1.3E-02 |
| REACTOME_PATHWAY | R-HSA-5357905 (Regulation of TNFR1 signaling) | 4 | 4.4 | 2.2E-03 | 2.7E-02 |
| **NF-kB signalling** | | | | | |
| KEGG_PATHWAY | Toll-like receptor signaling pathway | 14 | 15.6 | 1.4E-10 | 3.6E-09 |
| GOTERM_BP_DIRECT | positive regulation of I-kappaB kinase/NF-kappaB signaling | 12 | 13.3 | 8.6E-10 | 2.4E-07 |
| KEGG_PATHWAY | NF-kappa B signaling pathway | 12 | 13.3 | 3.0E-09 | 3.8E-08 |
| KEGG_PATHWAY | NOD-like receptor signaling pathway | 11 | 12.2 | 4.1E-10 | 7.9E-09 |
| GOTERM_BP_DIRECT | positive regulation of NF-kappaB transcription factor activity | 11 | 12.2 | 2.1E-09 | 4.8E-07 |
| KEGG_PATHWAY | RIG-I-like receptor signaling pathway | 11 | 12.2 | 4.9E-09 | 5.7E-08 |
| KEGG_PATHWAY | Cytosolic DNA-sensing pathway | 10 | 11.1 | 3.5E-08 | 3.0E-07 |
| REACTOME_PATHWAY | R-HSA-445989 (TAK1 activates NFkB by phosphorylation and activation of IKKs complex) | 8 | 8.9 | 2.6E-09 | 7.5E-07 |
| REACTOME_PATHWAY | R-HSA-168638 (NOD1/2 Signaling Pathway) | 8 | 8.9 | 1.0E-08 | 1.5E-06 |
| BIOCARTA | NF-kB Signaling Pathway | 7 | 7.8 | 1.2E-04 | 3.9E-03 |
| REACTOME_PATHWAY | R-HSA-1810476 (RIP-mediated NFkB activation via ZBP1) | 6 | 6.7 | 5.0E-07 | 4.8E-05 |
| REACTOME_PATHWAY | R-HSA-933542 (TRAF6 mediated NF-kB activation) | 6 | 6.7 | 1.0E-06 | 7.5E-05 |
| REACTOME_PATHWAY | R-HSA-2871837 (FCERI mediated NF-kB activation) | 6 | 6.7 | 4.5E-03 | 4.9E-02 |
| GOTERM_BP_DIRECT | nucleotide-binding oligomerization domain containing signaling pathway | 5 | 5.6 | 8.6E-06 | 5.4E-04 |
| GOTERM_BP_DIRECT | TRIF-dependent toll-like receptor signaling pathway | 5 | 5.6 | 1.4E-05 | 7.9E-04 |
| GOTERM_BP_DIRECT | I-kappaB kinase/NF-kappaB signaling | 5 | 5.6 | 2.9E-04 | 9.2E-03 |
| GOTERM_BP_DIRECT | NIK/NF-kappaB signaling | 5 | 5.6 | 4.1E-04 | 1.1E-02 |
| REACTOME_PATHWAY | R-HSA-5357956 (TNFR1-induced NFkappaB signaling pathway) | 4 | 4.4 | 1.7E-03 | 2.6E-02 |
| GOTERM_BP_DIRECT | negative regulation of NF-kappaB transcription factor activity | 4 | 4.4 | 6.3E-03 | 9.2E-02 |
| REACTOME_PATHWAY | R-HSA-5668541 (TNFR2 non-canonical NF-kB pathway) | 4 | 4.4 | 1.7E-02 | 1.3E-01 |
| **MAPK signalling** | | | | | |
| KEGG_PATHWAY | MAPK signaling pathway | 9 | 10.0 | 8.4E-03 | 2.3E-02 |
| GOTERM_BP_DIRECT | MAPK cascade | 7 | 7.8 | 2.7E-03 | 4.6E-02 |
| BIOCARTA | MAPKinase Signaling Pathway | 7 | 7.8 | 9.7E-02 | 3.8E-01 |
| GOTERM_BP_DIRECT | activation of MAPK activity | 5 | 5.6 | 2.5E-03 | 4.4E-02 |
| GOTERM_BP_DIRECT | positive regulation of MAPK cascade | 4 | 4.4 | 9.1E-03 | 1.2E-01 |
| REACTOME_PATHWAY | R-HSA-5673001 (RAF/MAP kinase cascade) | 4 | 4.4 | 6.4E-02 | 3.4E-01 |

### Supplementary Table S6

| Category | Term | Count | % | P-Value | Benjamini |
|---|---|---|---|---|---|
| TCR Activation | | | | | |
| KEGG_PATHWAY | T cell receptor signaling pathway | 8 | 15.1 | 3.2E-06 | 2.8E-05 |
| GOTERM_BP_DIRECT | T cell receptor signaling pathway | 7 | 13.2 | 6.2E-06 | 5.8E-04 |
| REACTOME_PATHWAY | R-HSA-202424 (Downstream TCR signaling) | 5 | 9.4 | 1.2E-03 | 2.2E-02 |
| BIOCARTA | T Cell Receptor Signaling Pathway | 4 | 7.5 | 4.9E-02 | 2.1E-01 |
| Apoptosis | | | | | |
| GOTERM_BP_DIRECT | apoptotic process | 14 | 26.4 | 1.2E-08 | 3.8E-06 |
| UP_KEYWORDS | Apoptosis | 13 | 24.5 | 6.9E-09 | 2.9E-07 |
| GOTERM_BP_DIRECT | regulation of apoptotic process | 9 | 17.0 | 2.7E-07 | 4.3E-05 |
| GOTERM_BP_DIRECT | positive regulation of apoptotic process | 7 | 13.2 | 3.2E-04 | 1.2E-02 |
| GOTERM_BP_DIRECT | negative regulation of apoptotic process | 7 | 13.2 | 2.7E-03 | 6.8E-02 |
| KEGG_PATHWAY | Apoptosis | 6 | 11.3 | 4.1E-05 | 2.7E-04 |
| GOTERM_BP_DIRECT | activation of cysteine-type endopeptidase activity involved in apoptotic process | 5 | 9.4 | 1.3E-04 | 5.6E-03 |
| BIOCARTA | Neuropeptides VIP and PACAP inhibit the apoptosis of activated T cells | 4 | 7.5 | 1.2E-02 | 7.5E-02 |
| BIOCARTA | Induction of apoptosis through DR3 and DR4/5 Death Receptors | 4 | 7.5 | 2.3E-02 | 1.2E-01 |
| TNF-alpha Signalling | | | | | |
| KEGG_PATHWAY | TNF signaling pathway | 11 | 20.8 | 7.5E-10 | 2.2E-08 |
| BIOCARTA | TNF/Stress Related Signaling | 8 | 15.1 | 1.7E-07 | 1.0E-05 |
| BIOCARTA | TNFR2 Signaling Pathway | 6 | 11.3 | 1.3E-05 | 4.0E-04 |
| GOTERM_BP_DIRECT | cellular response to tumor necrosis factor | 4 | 7.5 | 4.8E-03 | 9.9E-02 |
| GOTERM_BP_DIRECT | tumor necrosis factor-mediated signaling pathway | 4 | 7.5 | 5.8E-03 | 1.1E-01 |
| NF-KB Signalling | | | | | |
| KEGG_PATHWAY | NF-kappa B signaling pathway | 11 | 20.8 | 1.0E-10 | 1.5E-08 |
| KEGG_PATHWAY | Toll-like receptor signaling pathway | 11 | 20.8 | 7.5E-10 | 2.2E-08 |
| KEGG_PATHWAY | RIG-I-like receptor signaling pathway | 10 | 18.9 | 3.5E-10 | 1.7E-08 |
| GOTERM_BP_DIRECT | positive regulation of I-kappaB kinase/NF-kappaB signaling | 10 | 18.9 | 1.4E-09 | 1.3E-06 |
| GOTERM_BP_DIRECT | positive regulation of NF-kappaB transcription factor activity | 8 | 15.1 | 1.7E-07 | 3.1E-05 |
| KEGG_PATHWAY | NOD-like receptor signaling pathway | 9 | 17.0 | 1.4E-09 | 3.3E-08 |
| REACTOME_PATHWAY | R-HSA-168638 (NOD1/2 Signaling Pathway) | 7 | 13.2 | 1.2E-08 | 2.7E-06 |
| REACTOME_PATHWAY | R-HSA-933542 (TRAF6 mediated NF-kB activation) | 6 | 11.3 | 6.6E-08 | 7.5E-06 |
| REACTOME_PATHWAY | R-HSA-445989 (TAK1 activates NFkB by phosphorylation and activation of IKKs complex) | 6 | 11.3 | 2.2E-07 | 1.6E-05 |
| BIOCARTA | NF-kB Signaling Pathway | 6 | 11.3 | 4.8E-05 | 1.2E-03 |
| REACTOME_PATHWAY | R-HSA-1810476 (RIP-mediated NFkB activation via ZBP1) | 5 | 9.4 | 2.2E-06 | 1.0E-04 |
| REACTOME_PATHWAY | R-HSA-2871837 (FCERI mediated NF-kB activation) | 5 | 9.4 | 3.6E-03 | 4.7E-02 |
| GOTERM_BP_DIRECT | nucleotide-binding oligomerization domain containing signaling pathway | 4 | 7.5 | 6.1E-05 | 3.6E-03 |
| GOTERM_BP_DIRECT | toll-like receptor signaling pathway | 4 | 7.5 | 7.8E-05 | 4.0E-03 |
| GOTERM_BP_DIRECT | TRIF-dependent toll-like receptor signaling pathway | 4 | 7.5 | 8.7E-05 | 4.3E-03 |
| GOTERM_BP_DIRECT | NIK/NF-kappaB signaling | 4 | 7.5 | 1.1E-03 | 3.4E-02 |
| MAPK Signalling | | | | | |
| KEGG_PATHWAY | MAPK signaling pathway | 8 | 15.1 | 1.0E-03 | 4.6E-03 |
| BIOCARTA | MAPKinase Signaling Pathway | 7 | 13.2 | 4.9E-03 | 3.9E-02 |
| BIOCARTA | Human Cytomegalovirus and Map Kinase Pathways | 4 | 7.5 | 3.6E-03 | 3.3E-02 |
| GOTERM_BP_DIRECT | activation of MAPK activity | 4 | 7.5 | 4.4E-03 | 9.4E-02 |
| GOTERM_BP_DIRECT | MAPK cascade | 4 | 7.5 | 4.7E-02 | 4.3E-01 |
