## Supplementary Test for "An immunome perturbation is present in juvenile idiopathic arthritis patients who are in remission and will relapse upon anti-TNFα withdrawal"

**SUPPLEMENTARY ACKNOWLEDGMENTS AND AFFILIATIONS**

The authors would like to thank all the co-investigators from PRSCG of the clinical trial "Determining Predictors of Safe Discontinuation of Anti-TNF Treatment in JIA" (ID: NCT00792233) and are as follow; Authors (Daniel J. Lovell[1], Anne L. Johnson[1], Steven J. Spalding[2,17], Beth S. Gottlieb[3], Paula W. Morris[4], Yukiko Kimura[5], Karen Onel[17], Suzanne C. Li[5], Alexei A. Grom[1], Janalee Taylor[1], Hermine I. Brunner[1], Jennifer L. Huggins[1], James J. Nocton[7], Kathleen A. Haines[5], Barbara S. Edelheit[8], Michael Shishov[9], Lawrence K. Jung[10], Calvin B. Williams[7], Melissa S. Tesher[6], Denise M. Costanzo[2],[3], Lawrence S. Zemel[8], Jason A. Dare[4], Murray H. Passo[11], Kaleo C. Ede[9], Judyann C. Olson[7], Elaine A. Cassidy[12], Thomas A. Griffin[1], Linda Wagner-Weiner[6], Jennifer E. Weiss[5], Larry B. Vogler[13], Kelly A. Rouster-Stevens[13], Timothy Beukelman[14], Randy Q. Cron, [14], Daniel Kietz[12], Kenneth Schikler[15], Jay Mehta[16], Tracy V. Ting[1], James W. Verbsky[7], B. Anne Eberhard[3,5], Bin Huang[1], Chen Chen[1], Edward H. Giannini[1]) and Affiliations ([1]Cincinnati Children's Hospital Medical Center , Cincinnati, OH; [2]Cleveland Clinic Foundations, Cleveland, OH; [3]The Steven and Alexandra Cohen Children's Medical Center of New York, New Hyde Park, NY; [4]University of Arkansas for Medical Science, Little Rock, AR; [5]Hackensack University Medical Center, Joseph M. Sanzari Children's Hospital, Hackensack, NJ; [6]University of Chicago, Comer Children's Hospital, Chicago, IL; [7]Medical College of Wisconsin, Milwaukee, WI; [8]Connecticut Children's Medical Center, Hartford, CT; [9]Phoenix Children's Hospital, Phoenix, AZ; [10]Children's National Medical Center, Washington, DC; [11]Medical University of South Carolina, Charleston, SC; [12]Children's Hospital of Pittsburgh, Pittsburgh, PA; [13]Emory University School of Medicine, Atlanta, GA; [14]University of Alabama at Birmingham; [15]University of Louisville, Kosair Charities Pediatric Clinical Research Unit,

Louisville, KY; [16]Children's Hospital at Montefiore, Bronx, NY; [17] Hospital for Special Surgery, Weill Cornell Medicine, New York, NY).

### SUPPLEMENTARY METHODS

#### PBMC isolation and cryopreservation

The blood was drawn into anti-coagulation EDTA tubes and transported overnight to Sanford-Burnham institute and batch processed within the 16-20hrs window. PBMCs were isolated via density gradient centrifugation with Histopaque-1077 (Sigma-Aldrich) or Ficoll (GE Healthcare) according to the manufacturer's instructions. The cells were re-suspended in 90% v/v FBS, 10% v/v DMSO and frozen in liquid nitrogen for long-term storage.

#### PBMC interrogation by CyToF

PBMCs were stained with a T-cell focused panel of 37 heavy metal-conjugated antibodies (**online Supplementary Table S3**), as described previously[1] and subsequently analyzed by CyToF. Briefly, PBMCs were stimulated with or without phorbol 12-myristate 13-acetate (150 ng/ml, Sigma-Aldrich) and ionomycin (750 ng/ml, Sigma-aldrich) for 6 hrs, and blocked with secretory inhibitors, brefeldin A (1:1000, eBioscience) and monesin (1:1000, Biolegend) for the last 4 hrs, in 10% v/v human serum, 1% v/v PSG, RPMI at 37°C, 5% $CO_2$. The cells were then washed and stained with cell viability dye cisplatin (200 μM, Sigma-aldrich) for 5 min on ice. The cells were re-washed and each individual sample was barcoded with a unique combination of anti-CD45 conjugated with either heavy metal 89, 115, 141 or 167, as previously [2] for 25 min on ice. The barcoded cells were washed and stained with a surface antibody cocktail in 4% v/v heat-inactivated FBS, 2mM EDTA, 0.05% w/v sodium azide in pH 7.4 PBS for 30 mins on ice. The cells were again washed and re-suspended in fixation/permeabilization buffer (1:3, eBioscience) for 45 mins on ice. The permeabilized cells were then stained with an intra-antibody cocktail (1:10, permeabilisation buffer, eBioscience) for 45 mins on ice, before washing and staining with a DNA intercalator Ir-191/193 (1:2000 in 1.6% w/v paraformaldehye, Fluidigm) overnight at 4°C or for 20 mins on ice. Finally, the cells were washed and re-suspended with EQ™ Four Element Calibration beads (1:10, Fluidigm) in ultra-pure distilled water at $1x10^6$ cells /ml. The cell mixture was then loaded and acquired on a Helios mass cytometer (Fluidigm) calibrated with CyToF Tunning solution (Fluidigm). The output FCS files were randomized and normalized with the EQ™ Four Element Calibration beads against the entire run, as per the manufacturer's recommendations.

**CyToF data analysis with MarVis**

CyToF FCS data collection, conversion, randomisation, beads normalisation, concatenation were performed through the manufacturer's software (CyToF Software 6.7, Fluidigm). The pre-processed CyToF output FCS files (**online Supplementary Figure S8)** were de-barcoded manually (CD45- 89, 115, 141 or 167 metal) into individual samples in FlowJo (v.10.2). The $CD3^+CD4^+CD8^-CD14^-$ T cells were gated and exported for each individual sample. Batch effects were assessed across the 9 individual CyToF runs using an internal biological control, where one aliquot of the same healthy donor (PBMC) is used for each CyToF run (**online Supplementary Figure S1**). The exported CD4 T cells are normalised for equal cellular events. The CD4 T cells were then dimensionally reduced and clustered with MarVis (Multi-dimensional Automated Reduction and Visualisation) using Barnes Hut Stochastic Neighbour Embedding (SNE) non-linear dimensional reduction algorithm and k-means clustering algorithm, as previously described[1]. The default clustering parameters were set at perplexity of 30, and a minimum of $p < 0.0001$. The cells were then mapped on a 2-

dimensional t-distributed SNE scale based on the similarity score of their respective combination of markers, and categorized into nodes. The node phenotype was read using an R-script that compared the node marker intensity against the entire population of nodes in the histogram layout. The nodes were analyzed by Mann-Whitney U, two-tail test and a $p < 0.05$ was considered statistically significant. To ensure that the significant nodes obtained from clustering were relevant, we performed back-gating of the clustered CSV file and supervised gating of the original FCS files with FlowJo as validation.

**Cell sorting and culturing**

PBMCs (2 x$10^7$ cells/ml) were thawed and stained with CD3-AF700 (UCHT1, Biolegend), CD14-APC/H7 (MφP-9, BD Biosciences), CD4-BV605 (OKT4, Biolegend), CD45RA-PE/Dazzle (HI100, Biolegend), and CD45RO-FITC (UCHL1, Biolegend) for 20 mins on ice. Cell viability was determined by Sytox Red staining (1:1000, Thermofisher scientific). $CD3^+CD14^-CD4^+CD45RO^+CD45RA^-$ T cells were sorted on a FACS Aria II (BD Biosciences), and doublets and dead cells were excluded. The FCS sorting data was collected via the BD FACSDiva software. The cells were then seeded in a 96 well plate (4x$10^4$ cells per well) and incubated for 24 hrs with soluble tetrameric anti-CD3/CD28 (1:100, Stemcell) in 10% v/v human serum, 1% v/v PSG, RPMI at 37°C and 5% $CO_2$.

**mRNA purification and Nanostring screening**

mRNA was extracted from cells using an Arcturus Picture RNA Isolation Kit (Thermofisher scientific), according to the manufacturer's instructions. The washed and eluted RNA was amplified using an nCounter Low RNA Input Amplification Kit (Nanostring). First strand cDNA synthesis was performed with reverse transcriptase and RT primer mix at 42°C for 60 mins. Multiplexed target enrichment was then performed with gene-specific primers for 579 immunological and 15 reference control genes (nCounter Immunology panel V2, Nanostring) for eight cycles. The amplified RNA samples were hybridized with capture/reporter probes (nCounter Immunology panel v2, Nanostring) at 65°C for 16 hrs. Finally, the samples were captured onto nCounter chips using the prep station and read with a digital analyzer under maximum sensitivity (555 FOVs).

**Nanostring data analysis**

Nanostring mRNA data was collected through the nCounter Digitial analyser and exported as a RCC file. The RCC files were pre-processed with nSlover (v3, Nanostring) software provided by the manufacturer. Gene expression values were normalized to a recommended set of housekeeping genes designated within the chips. Statistical filtering was performed in nSlover (v3, Nanostring), with Welch's t test $p < 0.05$, and fold difference $\geq 1.5$. Statistically significant genes were exported to the Database for Annotation, Visualization and Integrated Discovery (DAVID, v6.8) website. Functional gene enrichment was performed with DAVID under the human background gene list. Genes from clusters of pathways that are significantly represented in DAVID were mapped and graphically represented with the Reactome database using Cytoscape (v3.5.1).

**Statistical analysis**

The non-parametric, non-paired, Mann-Whitney U, two tail statistical test was performed in MarVis clustering and manual gating of individual cellular subsets in FlowJo; * $p < 0.05$, ** $p < 0.01$, *** $p < 0.001$ and **** $p < 0.0001$. For the comparison of percentage of cells in nodes generated through clustering, a correction is performed for different numbers of subjects, and reflected as relative percentage. For the testing of paired cellular subsets within each individual, we performed the non-parametric, Wilcoxon matched pairs signed rank two tail test; * $p < 0.05$, ** $p < 0.01$, *** $p < 0.001$ and **** $p < 0.0001$. For correlation analysis, the Pearson correlation two tail test was used. For group analysis across ages, Kruskal-Wallis test was used. The Mann-Whitney U, Wilcoxon test, Pearson correlation, Kruskal-Wallis, and ROC statistical testing were performed through GraphPad Prism (version 7.0e). Nanostring differential gene expression comparison was performed with the Welch's t test as designated in the nSlover statistical software provided by manufacturer.

### SUPPLEMENTARY METHODS REFERENCES


1. Chew V, Lee YH, Pan L, et al. Immune activation underlies a sustained clinical response to Yttrium-90 radioembolisation in hepatocellular carcinoma. Gut. 2018.
2. Lai L, Ong R, Li J, et al. A CD45-based barcoding approach to multiplex mass-cytometry (CyTOF). Cytometry A. 2015;87(4):369-74.


### SUPPLEMENTARY FIGURE AND TABLE LEGENDS

**Supplementary Figure S1**. **Internal biological controls for accessing batch effects of individual CyToF runs.** To access staining variations across the batch runs (1-9), one aliquot of the same healthy donor was run for each individual CyToF run. The staining intensity for every marker is depicted against individual runs for the total PBMC population.

**Supplementary Figure S2. Density plots for marker expression in t-SNE map.** Dimensional reduction with t-SNE of $CD3^{+}CD4^{+}$ T cells from relapse and remission patients (n=20) using MarVis. Density expression maps depicting the distribution and expression of cells with the 31 functional markers as shown.

**Supplementary Figure S3. Gating strategy for cellular subsets.** The gating strategy is shown for the relevant cellular subsets described in accordance with the following markers CD45RA, TNFα, IFNY, IL-6, CD152, PD1, CD25, FoxP3.

**Supplementary Figure S4. Sorting strategy for $CD3^+CD4^+CD14^-CD45RO^+CD45RA^-$ cells.** PBMCs were thawed and stained with the respective antibodies and sorted for $CD3^+CD4^+CD45RA^-CD45RO^+$ T cells using a FACS Aria II. Doublets and dead cells were excluded.

**Supplementary Figure S5. Persistence of genes in relapse or remission individuals despite anti-TNFα therapy.** Equal numbers of sorted $CD3^+CD4^+CD14^-CD45RO^+CD45RA^-$ T cells from relapse (n=6, $T_o/T_{end}$), remission (n=6, $T_o$ / $T_{end}$) patients with JIA and healthy pediatric controls (n=3) were stimulated for 24hrs with anti-CD3/CD28, and subjected to mRNA analysis using the Nanostring Immunology V2 panel. Heatmap depicting genes significantly increased in (**A**) relapse or (**B**) remission ($T_o$ / $T_{end}$) patients with JIA compared to healthy controls ($p < 0.05$, fold difference ±1.5). Genes highlighted in blue are enriched in remission individuals; genes highlighted in red have been previously described in GWAS JIA studies.

**Supplementary Figure S6. Persistence of genes in JIA patients susceptible to anti-TNFα therapy from pre to post treatment.** Equal numbers of sorted $CD3^+CD4^+CD14^-$ $CD45RO^+CD45RA^-$ T cells from four paired patients with JIA pre (active) or post (recent onset clinical inactivity) anti-TNFα therapy, and three healthy paediatric controls were stimulated for 24hrs with anti-CD3/CD28, and subjected to mRNA analysis with Nanostring Immunology V2 panel. Heatmap showing differential expressed genes (D.E.Gs) significantly

($p < 0.05$, fold difference ±1.5 ) increased in patients with JIA compared to healthy controls without CD3/CD28 stimulation.

**Supplementary Figure S7. Pathway analysis of persistent genes in JIA patients susceptible to anti-TNFα therapy from pre to post treatment.** Equal numbers of sorted $CD3^+CD4^+CD14^-CD45RO^+CD45RA^-$ T cells from four paired patients with JIA pre (active) or post (recent onset clinical inactivity) anti-TNFα therapy, and three healthy paediatric controls were stimulated for 24hrs with anti-CD3/CD28. Genes enriched in patients with JIA were exported to DAVID for functional gene-set enrichment, and gene associations were constructed with Cytoscape using the Reactome database. Five major pathways (**A**) TCR activation, (**B**) TNFα signalling, (**C**) NF-kB signalling, (**D**) Apoptosis, (**E**) MAPK signalling were dysregulated in patients with JIA compared to healthy controls.

**Supplementary Figure S8. Pre-processing of CyToF FCS files and gating for $CD3^+CD4^+CD14^-CD8^-$ T cells.** CyToF FCS files were pre-processed for singlets, live cells, stringent EQ-beads removal and CD45 debarcoded into individual sample files. Each sample file is subsequently gated for $CD3^+CD4^+CD8^-CD14^-$ T cells as shown.

**Supplementary Table S1**. **Demographics and medication history of the JIA patients withdrawn from therapy and healthy controls.** Tables depicting the usage of samples across the experiments and the demographic summary table. Patients with JIA were administered anti-TNFα medication (etanercept, adalimumab or infliximab) with or without concurrent methotrexate (MTX) combination, for at least 6 months, and proven to be in inactive disease (Wallace criteria) before anti-TNFα withdrawal. Each JIA patient was scored as either relapse or remission. Healthy non-disease controls with no inflammatory diseases were recruited from day surgeries. $T_o$ : PBMCs obtained prior to therapy withdrawal; $T_{end}$ : PBMCs obtained by the end of 8 months after therapy withdrawal.

**Supplementary Table S2**. **Disease and medication history of patients with active JIA.** $CD3^+$ $CD4^+$ $CD14^-$ $CD45RA^-$ $CD45RO^+$ T cells were sorted from patients with active JIA paired for pre (treatment naive) and post (recent onset clinical remission) anti-TNFα therapy, and then subjected to mRNA analysis with Nanostring.

**Supplementary Table S3. CyToF antibody panel used for staining PBMCs from patients with JIA and controls.** The list of antibodies and heavy metals conjugated, with vendor catalogue number is shown.

**Supplementary Table S4. DAVID functional gene-set enrichment of genes enriched in relapse ($T_o$ / $T_{end}$) individuals.** DAVID functional gene-set enrichment was performed for genes enriched (DEGs $p < 0.05$, fold difference > 1.5) in relapse ($T_o$ / $T_{end}$) individuals compared to healthy controls, with default setting against a human background. Pathways implicated are tabulated for gene counts ≥ 4.

**Supplementary Table S5. DAVID functional gene-set enrichment of genes enriched in remission ($T_o$ / $T_{end}$) individuals.** DAVID functional gene-set enrichment was performed for genes enriched (DEGs $p < 0.05$, fold difference > 1.5) in remission ($T_o$ / $T_{end}$) individuals compared to healthy controls, with default setting against a human background. Pathways implicated are tabulated for gene counts ≥ 4.

**Supplementary Table S6. DAVID functional gene-set enrichment of genes enriched in JIA patients susceptible to anti-TNFα therapy from pre to post treatment.** DAVID functional gene-set enrichment was performed for genes enriched (DEGs $p < 0.05$, fold difference > 1.5) in JIA pre (active) or post (recent onset clinical inactivity) anti-TNFα therapy, and three healthy paediatric controls, with default setting against a human background. Pathways implicated are tabulated for gene counts ≥ 4.
